# A tumour-host feed-forward loop contributes to malignant growth in chromosomal instability-induced epithelial tumours

**DOI:** 10.1101/2025.09.24.678048

**Authors:** Kaustuv Ghosh, Aishwarya Kunchur, Marco Milán

**Affiliations:** Institute for Research in Biomedicine (IRB Barcelona), The Barcelona Institute of Science and Technology, Baldiri Reixac, 10, 08028 Barcelona, Spain; Institució Catalana de Recerca i Estudis Avançats (ICREA) Pg. Lluís Companys 23, 08010 Barcelona, Spain

**Keywords:** aneuploidy, senescence, SASP, super-competition, hippo signalling

## Abstract

Chromosomal instability (CIN), characterized by frequent changes in chromosome number and structure, is common in human carcinomas and often leads to aneuploidy. Drosophila has been instrumental in demonstrating that CIN can promote tumour growth and malignancy through aneuploidy-induced senescence, a state marked by cell-cycle arrest and high secretory activity. Despite extensive chromosomal heterogeneity, we show that these cells share a distinct transcriptional program, with most responses to aneuploidy and senescence regulated at the transcriptional level. Nearly 10% of the most upregulated genes encode secreted proteins of the senescence-associated secretory phenotype. Five of these proteins act additively, locally or systemically, to block proliferation and induce cell death in neighbouring tissues. This non-autonomo*us cell death feeds back to the tumour to enhance its growth, resembling super-competition and providing insight into tumour–host interactions relevant to human cancer.

**Teaser:** Aneuploidy-induced senescent cells activate the pro-survival Hippo-Yorkie pathway and the SASP drives tumour host interactions

## Introduction

Chromosomal instability (CIN), an increased rate of errors in chromosome segregation during cell division, is one of the most common forms of genomic instability in cancer. It is prevalent in most solid tumours and is closely associated with chemotherapy resistance, reduced patient survival, and metastasis (*1–6*). Traditionally, CIN has been understood to be a driver of genetic diversity within tumours, mainly through the gradual gain of chromosomes that carry oncogenes and the loss of those containing tumour suppressor genes. However, beyond these slow, progressive changes, CIN can also cause sudden and dramatic alterations in the genome. For example, mis-segregated chromosomes can end up in micronuclei, where they often experience extensive DNA damage. This process, known as chromothripsis, leads to the fragmentation and chaotic rearrangement of chromosomes (*7*). In addition to fuelling cancer progression through both gradual and abrupt mechanisms, CIN also causes cytosolic double-stranded DNA, likely originating from ruptured micronuclei. This DNA is detected by cellular sensors, such as cGAS, which normally activate immune responses. However, cancer cells can exploit this pathway to reshape the tumour microenvironment in a way that suppresses immune activity and promotes metastasis (*8–10*).

Although low levels of CIN are widely recognized as a driver of cancer, excessive levels can be harmful, leading to cell death and the suppression of tumour growth (*11–14*). In normal cells, CIN often results in aneuploidy—an abnormal number of chromosomes—which can disrupt cell function (*15*). Studies in yeast and human cells have shown that this imbalance in the number of chromosomes alters protein ratios and interferes with key cellular processes like DNA replication, mitosis, and metabolism (*16–18*). One outcome of such metabolic disruption is the production of reactive oxygen species (ROS). Studies in *Drosophila* and human epithelial cells subjected to CIN have shown that cells with aneuploid karyotypes that affect a large proportion of the genome typically enter a state of senescence, where they permanently stop dividing and begin releasing inflammatory molecules (*19–22*). These molecules help the mammalian immune system identify and eliminate abnormal cells (*23*). Research in *Drosophila* has further revealed that aneuploidy-induced senescence promotes tumour development. The fly genome is organized into four chromosomes [two autosomes, a sex chromosome pair (X and Y), and a tiny dot-like autosome]. In epithelial tissues subjected to CIN, random chromosome mis-segregation events generate cells with whole-chromosome aneuploidies (thus impacting a large proportion of the genome) that are expelled from tissues as a result of gene dosage and proteome imbalances, proteostasis failure, and mitochondrial dysfunction (*19*, *24*, *25*). Once extruded, these cells undergo apoptosis as a result of ROS-driven activation of JNK and JAK/STAT signalling pathways (*19*, *24*, *26*, *27*). When fly aneuploid cells are maintained in the tissue by apoptosis inhibition, they often undergo a JNK-driven transition to a mesenchymal-like state and enter a form of senescence marked by G2 cell cycle arrest (*19*, *28*). Key molecules expressed by senescent cells include the *Drosophila* orthologs of Wnt and IL-6 [Wingless (Wg) and Upd3, respectively], which act on the epithelial tissue subjected to CIN to drive tumour growth, to enhance cell motility of aneuploidy-induced delaminated cells, and even affect systemic hormonal signals that prevent developmental maturation and cause organismal death (*24*, *29–31*).

In this study, we used the *Drosophila* epithelial model of CIN-induced tumorigenesis to characterize the transcriptional profile of aneuploidy-induced senescent cells. Despite the highly heterogeneous chromosome content of these cells, we unravel a distinct transcriptional program. This unique transcriptional program presents evidence that most cellular responses to aneuploidy and senescence, including cell cycle arrest, activation of autophagy, reorganization of the actin cytoskeleton, and a highly secretory phenotype, are regulated at the transcriptional level. In addition to the signalling pathways involved in driving all these cellular responses, we unravel a pro-survival function of the Hippo-Yorkie signalling pathway in aneuploid senescent cells. Besides the secreted proteins well known to promote tumour growth, metastasis, and malignancy (*24*, *29–31*), we uncovered a role of dIlp8 and ImpL2 (fly orthologs of Relaxin and IGBP7, respectively (*32–34*)) in blocking proliferative growth of both wildtype and tumour cells, and identified a role for cytokines Upd1 and Upd3 and the Drosophila Tumour Necrosis Factor (TNF-α) homologue Eiger in non-autonomously driving cell death in neighbouring wild type cells. Most importantly, we present evidence that this non-autonomous cell death feeds back to the tumour to enhance its growth, thus paralleling the phenomenon of super-competition in cancer, an aggressive form of cell competition where tumour cells actively eliminate neighbouring, healthy wild-type cells to expand its territory (*35–37*). Overall, our data highlight potential therapeutic strategies to prevent CIN-driven tumour development by targeting aneuploidy-induced senescent cells and disrupting the relevant interactions with the tumour host.

## Results

### Cellular responses to aneuploidy and senescence are reflected at the transcriptional level

We used the wing primordium of *Drosophila*—a highly proliferative epithelial monolayer that forms a two-sided epithelial sac (Figure 1A)—as a model system to characterize the transcriptional profile of cells subjected to CIN. CIN, an increased rate of changes in chromosome structure and number, was induced by depletion of the spindle assembly checkpoint (SAC) gene *rod* (*rough deal*) using an RNAi form expressed with the GAL4/UAS system (*24*, *26*). The *engrailed-gal4* driver, which is restricted to the posterior (P) compartment of the wing primordium (Figure 1A), was used to limit the induction of CIN to a defined cell population. One of the most attractive features of this model of CIN is that it facilitates the identification of three well-defined cell populations with distinct behaviours (Figure 1B). On one hand, if apoptosis is blocked by the expression of p35 (a baculovirus protein that binds and represses effector caspases Dcp-1 and Drice (*38*)), those cells that become aneuploid - as a consequence of random chromosome missegregation events in mitosis caused by *rod* depletion - delaminate and become cell cycle-arrested, motile, and highly secretory. These cells can then be tracked with MMP1-GFP, which is a GFP-based transcriptional reporter of JNK (*39*) (Figure 1B, green cells). On the other hand, the epithelial tissue subjected to CIN (the P compartment), which consists of cells that remain largely euploid, enters a hyperplastic program that relies on high mitotic activity and unlimited growth potential (Figure 1B, cells located apically to the green cells). Finally, the anterior (A) compartment of these wing primordia (Figure 1B, cells labelled by the Ci marker in red), which are not subject to CIN and whose cells are euploid, will be treated as the neighbouring cell population. The transcriptional profile of these three cell populations (CIN, aneuploid cells, and neighbouring cell population) was compared to the transcriptional profile of control wing primordia expressing p35 under the control of the *engrailed-gal4* driver. Four distinct biological replicates were analysed. Principal Component Analysis (PCA) revealed significant transcriptomic differences between the five populations, showing strict clustering based on their respective traits (Figure 1C). Interestingly, the aneuploid cell population cluster presented the highest level of difference when compared to the compartment of origin [P(CIN)] or the control tissue [P(Control)]. Consistent with this observation, the aneuploid population showed a higher number of upregulated and downregulated genes than the CIN epithelium (Figure 1D, F, see also Figure S2A). However, there was a certain degree of overlap in the type of genes that changed in these two populations when compared to control tissues. We thus analysed the transcriptional profile that was unique to aneuploid cells. Remarkably, many of the previously observed cellular responses to aneuploidy were reflected at the transcriptional level, as shown by gene set enrichment analysis (GSEA, Figure 1E, Figure S1A, and Table S1). First, high levels of proteotoxic stress caused by unbalanced karyotypes were reflected by strong upregulation of genes involved in autophagy (e.g., *atg4a*, *atg8a*, *atg9*, *atg17*, and *atg18b*) and lysosome function (e.g., *Lamp1*, *Vps13D*, *Vps16A, Vamp7,* and *lqf*). Second, consistent with the activation of the JNK stress signalling pathway as a consequence of mitochondrial dysfunction and ROS (*19*, *27*), many of its target genes (e.g., *mmp1*, *ets21C*, and *scaf*) were strongly upregulated in aneuploid cells (Figure 1E, 2A, and Figure S1A). Third, the ability of aneuploid cells to become motile through modulation of the actin cytoskeleton and a non-apoptotic role of caspases (*28*, *31*) was reflected by strong upregulation of genes involved in these two processes, as well as in cell motility (e.g., *sqh*, *cortactin*, *rok*, *fak*, *Arpc2-5, hid*, *rpr*, *Dronc*, *Drice*, and *Dcp-1;* Figures 1E, 2A, and Figure S1A). Finally, entry into a senescence-like state ^1^ was also reflected at the transcriptional level, as shown by strong downregulation of genes involved in cell cycle progression (e.g., *stg*, *cdk1*, *cycA*, and *cycB*) and DNA replication (e.g., *DNA polymerase*, *Orc* and *Mcm* complexes, Figure 1E and 2B), and the clear upregulation of genes involved in secretion (e.g., *Tango 5, 11, Sec 5, 6, 33, 24CD, and Rab 1, 2, 6, 7, 10,* and *35*) and encoding for secreted proteins (Figure 1E, 2A, and S1A). Indeed, 10% of the most upregulated genes encoded for secreted proteins (Table 1 and S2). These included the systemic signals Upd3 cytokine (involved in CIN-induced invasion and developmental delay (*30*, *31*)), Dilp8 relaxin (mediating developmental delay (*30*, *31*)), Pvf1 (PDGF- and VEGF-related factor 1 involved in driving organ wasting (*40*)) and the mitogenic signals Wg and Wnt6 (*29*), many of them are known targets of JNK. We confirmed the highly secretory phenotype of senescent cells by using three fluorescent reporters to mark the enlargement of the Golgi and the endoplasmic reticulum (ER) compartment, and an increase in the number of exosomes (Figure 2C). The transcriptional profile of aneuploid cells that was shared with CIN comprised genes involved in the activation of ERK and JAK/STAT signalling, previously shown to contribute to various aspects of CIN-induced tumorigenesis, such as growth and invasiveness (*28*, *31*) (Figure 1G and S2B, C). In CIN cells, we observed unique upregulation of the genes involved in DNA damage, consistent with the high propensity of dividing cells subjected to SAC depletion to accumulate lagging or broken chromosomes (Figure S2B, C). Overall, these results indicate that the cellular responses to aneuploidy and senescence are reflected at the transcriptional level, and they reinforce the highly secretory phenotype of aneuploidy-induced senescent cells.

**Figure 1.**
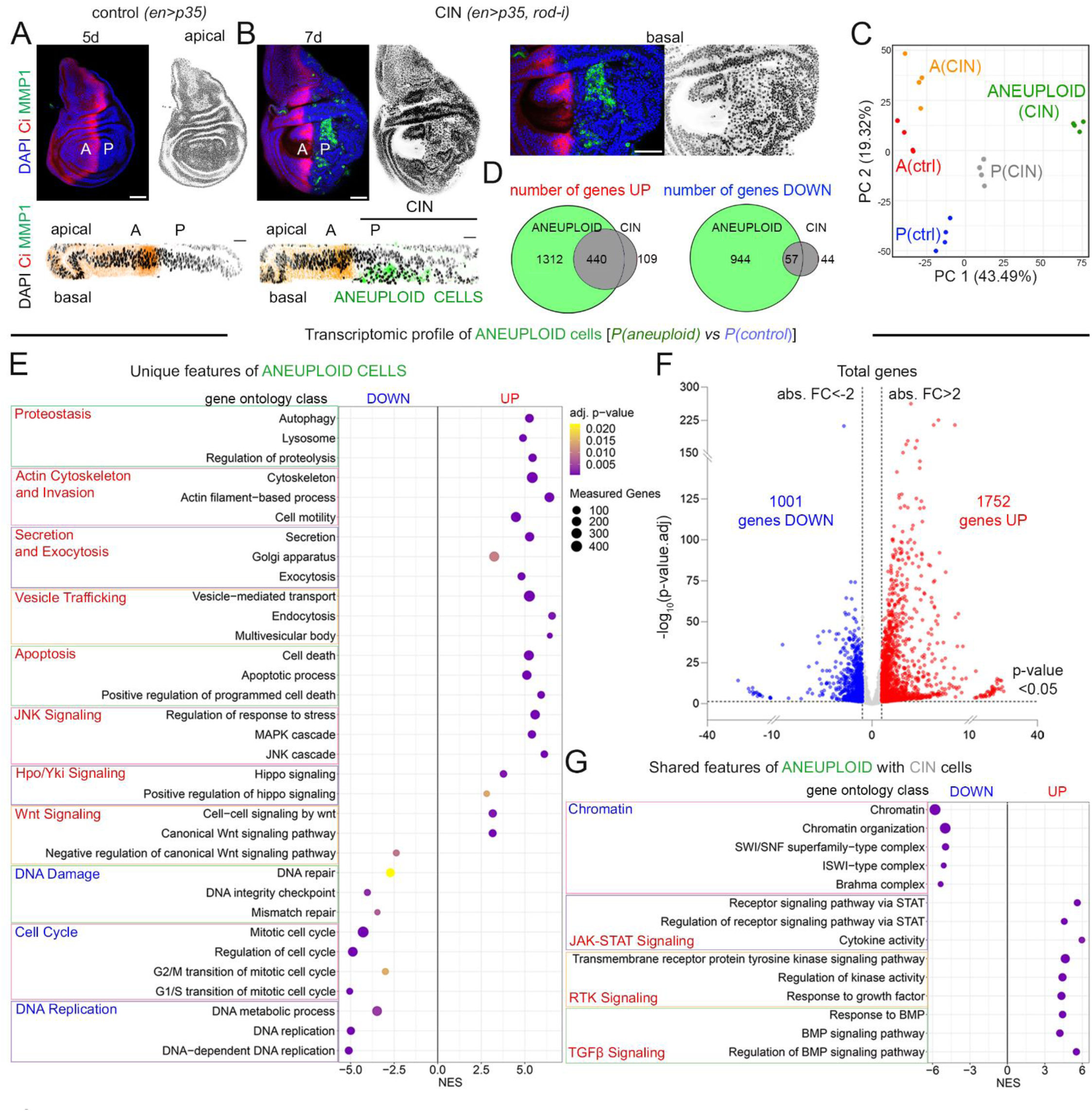
A distinct transcriptomic profile of aneuploidy-induced senescent cells. **(A, B)** Control wing discs (**A**) and wing discs subjected to CIN (**B**), stained for DAPI (blue or black), Ci (red, to label the A compartment), and MMP1 (green, to label the aneuploid cells) and expressing the indicated transgenes in the P compartment under the control of the *en-gal4* driver. Scale bars, 50 µm. Days of dissection after egg laying are as indicated. **(C)** Principal Component (PC) analysis showing the transcriptomic differences between the 5 different populations: A (Red) and P (Blue) cells of control wing discs, and aneuploid (green), A (orange) and P (gray) cells of wing discs subjected to CIN. **(D)** Venn diagram representing the number of genes up- or down-regulated uniquely in aneuploid cells (green), and proliferating cells subjected to CIN cells or in both types of cells (gray). **(E, G)** Bubble plots representing Gene Ontology (GO) enrichment analyses of aneuploid (**E**) and aneuploid and proliferating cells (**G**) when compared to P cells of control discs. Size of the bubble represents the number of measured genes, and the colour scale represents the associated p-value. **(F)** Volcano plot representing all the genes that are differentially up- (red dots, FC>2 and p-value<0.05) or down-regulated (blue dots FC<-2 and p-value<0.05), or not differentially expressed (gray dots, p-value>0.05 or 2>FC>-2) in aneuploid cells when compared to P cells of control discs. See also Figure S1, S2, and Table S1.

**Figure 2.**
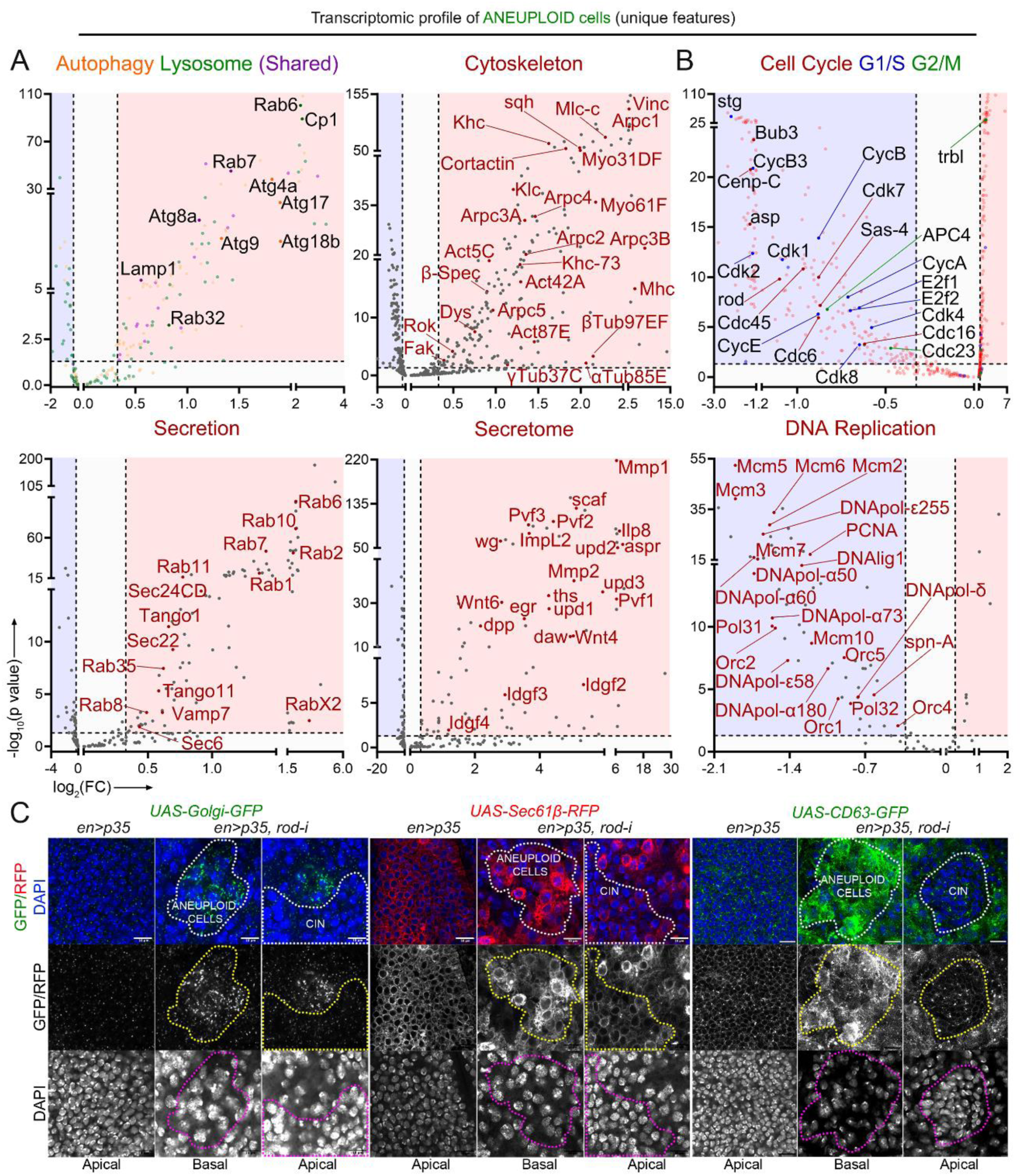
The transcriptome of aneuploidy-induced senescent cells points to a heightened secretory potential. **(A, B)** Volcano plot representing GO class-specific genes that are differentially upregulated (red area, FC>2 and p-value<0.05), downregulated (blue area, FC<-2 and p-value<0.05), or not differentially expressed (white area, p-value>0.05 or 2>FC>-2) in aneuploid cells when compared to P cells of control discs. GO classes (A): Autophagy - GO0006914, Lysosome - GO0005764, Cytoskeleton - GO0005856, Secretion - GO0046903, Secretome - Custom Gene Set. GO classes (B): G1/S Transition - GO0000082, G2/M Transition - GO0000086, Regulation of cell cycle - GO0051726, and DNA Replication - GO0006260. Genes corresponding to each of the GO classes have a distinct colour code. **(C)** High magnification images of wing discs expressing the indicated transgenes under the control of the *en-gal4* driver and stained to visualize differential expression of fluorescent reporters (green, red, or white) and DAPI (blue and white). Apical and basal sections are shown. Aneuploid and CIN cells are marked by a dashed line. Scale bars, 10 µm. See also Tables S1 and S2.

**Table 1.** The secretome of aneuploidy-induced senescent cells. Table representing the 40 most differentially upregulated genes (comparing the transcriptome of the aneuploidy-induced senescent cells with control cells) which are annotated as genes encoding for secreted proteins, along with their vertebrate homolog, adjusted p. values and shrunken fold-changes. The genes labelled in red are those which have been mentioned in the current work. A manually curated custom gene set, consisting of 261 known or predicted secreted proteins as inferred from FlyBase is provided in Table S2. The transcriptome of the aneuploidy-induced senescent cells was compared to posterior wild-type cells.

| Symbol | Gene name | Vertebrate homolog | Significance (p.Value) | Fold Change |
| --- | --- | --- | --- | --- |
| RYa | Ryamide | NPY family | 3,25E-08 | 21721233,75 |
| NLaz | Neural Lazarillo | ApoD and RBP4 | 1,07E-35 | 681,212 |
| aspr | asperous | DNER and EDIL3 | 1,32E-55 | 491,381 |
| Ilp8 | Insulin-like peptide 8 | Relaxin | 5,46E-81 | 198,57 |
| upd2 | unpaired 2 | Leptin | 3,26E-49 | 110,948 |
| nec | necrotic | SERPIN | 1,02E-75 | 89,581 |
| PGRP-SA | Peptidoglycan recognition protein SA | PGLYRP3 | 2,88E-59 | 88,703 |
| Mmp1 | Matrix metalloproteinase 1 | MMP | 2,30E-215 | 83,782 |
| Pvf1 | PDGF- and VEGF-related factor 1 | PDGF and VEGF | 4,28E-31 | 75,409 |
| Tig | Tiggrin | RGD-containing integrin ligand | 1,96E-26 | 73,827 |
| Sp7 | Serine protease 7 | Serine Proteases | 5,87E-85 | 65,623 |
| upd3 | unpaired 3 | Il-6 | 3,28E-34 | 56,97 |
| CG30280 | Unannotated | MFAP4 | 8,16E-05 | 43,886 |
| sona | sol narae | ADAMTS | 2,08E-120 | 41,327 |
| tnc | tenectin | Mucin-type protein with RGD motif | 4,07E-17 | 37,48 |
| scaf | scarface | Serine Protease | 2,42E-124 | 34,526 |
| Mmp2 | Matrix metalloproteinase 2 | MMP | 3,28E-39 | 31,565 |
| frm | farmer | FLG | 1,72E-144 | 30,547 |
| Wnt4 | Wnt oncogene analog 4 | WNT | 1,01E-14 | 28,179 |
| ldgf2 | Imaginal disc growth factor 2 | CHI3L1, CHI3L2, and OVGP1 | 5,07E-07 | 26,81 |
| daw | dawdle | NODAL | 1,84E-14 | 26,357 |
| Spn47C | Serpin 47C | SERPIN | 4,32E-71 | 23,661 |
| Pvf2 | PDGF- and VEGF-related factor 2 | PDGF and VEGF | 5,62E-99 | 21,107 |
| ths | thisbe | FGF8 | 1,46E-32 | 18,374 |
| CG31869 | Unannotated | KCP | 2,31E-86 | 18,285 |
| upd1 | unpaired 1 | Il-6 | 1,01E-26 | 18,278 |
| mtg | mind the gap | Lectin | 3,46E-90 | 13,673 |
| ImpL2 | Ecdysone-inducible gene L2 | IGFBP7 | 1,43E-76 | 12,527 |
| Pvf3 | PDGF- and VEGF-related factor 3 | PDGF and VEGF | 1,21E-92 | 12,395 |
| CecB | Cecropin B | Similar to Antimicrobial peptides | 1,15E-03 | 12,284 |
| spz | spatzle | Neurotrophin | 7,22E-09 | 11,326 |
| CG9095 | Unannotated | Secretory phospholipase A2 | 3,11E-19 | 10,982 |
| egr | eiger | TNF | 2,52E-22 | 10,773 |
| fon | fondue | NCOA5 and NUP98 | 3,01E-08 | 9,755 |
| l(1)G0193 | orion | Similar to Chemokines | 2,25E-56 | 8,84 |
| Dro | Drosocin | Similar to Antimicrobial peptides | 1,43E-03 | 8,007 |
| Spn55B | Serpin 55B | SERPIN | 1,94E-64 | 7,591 |
| Wnt6 | Wnt oncogene analog 6 | WNT | 1,32E-29 | 6,846 |
| wg | wingless | WNT | 5,12E-62 | 6,798 |
| Cp15 | Chorion protein 15 | No homolog identified | 4,48E-04 | 6,417 |

### A survival role of hippo/Yorkie in CIN-induced senescent cells

In CIN-induced tumours, the ability of aneuploid cells to become motile relies on modulation of the actin cytoskeleton and a non-apoptotic role of caspases (*28*, *31*). Consistent with this, activation of the apoptotic pathway in CIN tissues (monitored with cDcp1 antibodies, Figure 3D, E) was not followed by apoptosis execution (monitored by TUNEL staining, Figure 3C). Activation of the apoptotic pathway in aneuploidy-induced senescent cells was reflected by strong upregulation of genes involved in apoptosis, including the pro-apoptotic genes *hid* and *reaper*, the initiator caspase *Dronc,* and the effector caspases *Drice* and *Dcp-1 (*Figure 3A). The increase in *hid* expression was confirmed by a *hid-lacZ* reporter (Figure 3B). Interestingly, the activity levels of effector Caspases, monitored by the GC3Ai activity reporter (*41*), were relatively high in aneuploidy-induced senescent cells when compared to control tissues (Figure 3B) despite the expression of the p35 baculovirus protein, a potent inhibitor of these Caspases (*38*). However, there were no pyknotic nuclei in the tissue suggesting that apoptosis execution is indeed actively blocked in these cells. In our search for candidate pathways with well-known pro-survival activity, we observed that many elements of the Hippo/Yorkie signalling cascade, including *crumbs* (*crb*), *fat* (*ft*), *Merlin* (*Mer*), *Ajuba* (*jub*), *kibra*, *hippo* (*hpo*), and *yorkie* (*yki*), were transcriptionally activated in aneuploidy-induced senescent cells (Figure 3A). The activation of Yorkie in these cells was confirmed with the use of GFP-protein traps of Ajuba (Jub-GFP, an upstream negative regulator of Hippo (*42*)) and Yorkie (Figure 3C, F), transcriptional reporter and GFP-protein trap of *Diap1* (an E3 ubiquitin ligase regulated by Yorkie and involved in inhibiting Dronc, (*43*), Figure 3E and S3), and transcriptional reporters of *bantam* (a miRNA regulated by Yorkie and involved in inhibiting *hid* (*44*, *45*)), and *expanded* [an upstream regulator of the Hippo pathway and Yorkie target (*46*), Figure 3D, S3]. Yorkie has been previously shown to be activated by JNK (*47*), which is highly active in CIN-induced aneuploid cells and plays a pivotal role in defining their senescent state (*19*). Consistently, the expression levels of Yorkie targets *Diap1* and *bantam* were significantly reduced upon blockage of the JNK pathway by expression of a dominant negative version of JNK (Bsk in *Drosophila* (*48*), Figure 3D, E, quantified in Figure 3D’, E’). Given the well-known anti-apoptotic role of the Yorkie pathway, we addressed whether Yorkie depletion had any impact on the survival of CIN tissues. To this end, and based on the developmental role of Yorkie in regulating growth and survival of epithelial tissues (*49*), we searched for a mild *UAS-yorkie-RNAi* transgene. Interestingly, depletion of Yorkie (with the use of this mild transgene) did not have a major effect on the size or survival of p35-overexpressing tissues but did have a mild effect on the size and survival of otherwise wild-type tissues (Figure 4A, C, left panels, and quantified in Figure 4B, D). In contrast, the use of this transgene induced a strong negative impact on the size of CIN tissue, and this reduction was accompanied by a significant increase in the number of apoptotic cells (Figure 4A, C, right panels, and quantified in Figure 4B, D). In experimental conditions where apoptosis was not blocked, the CIN tissue collapsed and was transformed into isolated islands of dying cell populations (Figure 4C, right panel). Taken together, these results indicate that the Hippo/Yorkie pathway acts as an intrinsic anti-apoptotic pathway that contributes to dampening the deleterious effects of CIN and maintaining senescent cells alive despite the activation of the apoptotic pathway. Consistent with the proposal of this intrinsic anti-apoptotic pathway, we observed that by halving the doses of the pro-apoptotic genes *reaper*, *hid* and *grim* (in *Df(H99)/+* larvae), the levels of cell death observed in CIN tissues were reduced (Figure 4E, and quantified in Figure 4I), and aneuploid cells, labelled by the expression of MMP1, were maintained in the tissue even in the absence of p35 (Figure 4E and F). These cells were negative for apoptotic markers (Figure 4G) and were not the result of loss of Df(H99) heterozygosity caused by CIN, as they expressed the *bantam-lacZ* reporter located in the homologous chromosome (Figure 4H). Thus, aneuploid cells were not maintained in the tissue in this experimental setting as a trivial consequence of inheriting the two chromosomes bearing the *Df(H99)* deficiency. Most interestingly, under these circumstances, the tissue also overgrew (Figure 4E, F, and quantified in Figure 4I).

**Figure 3.**
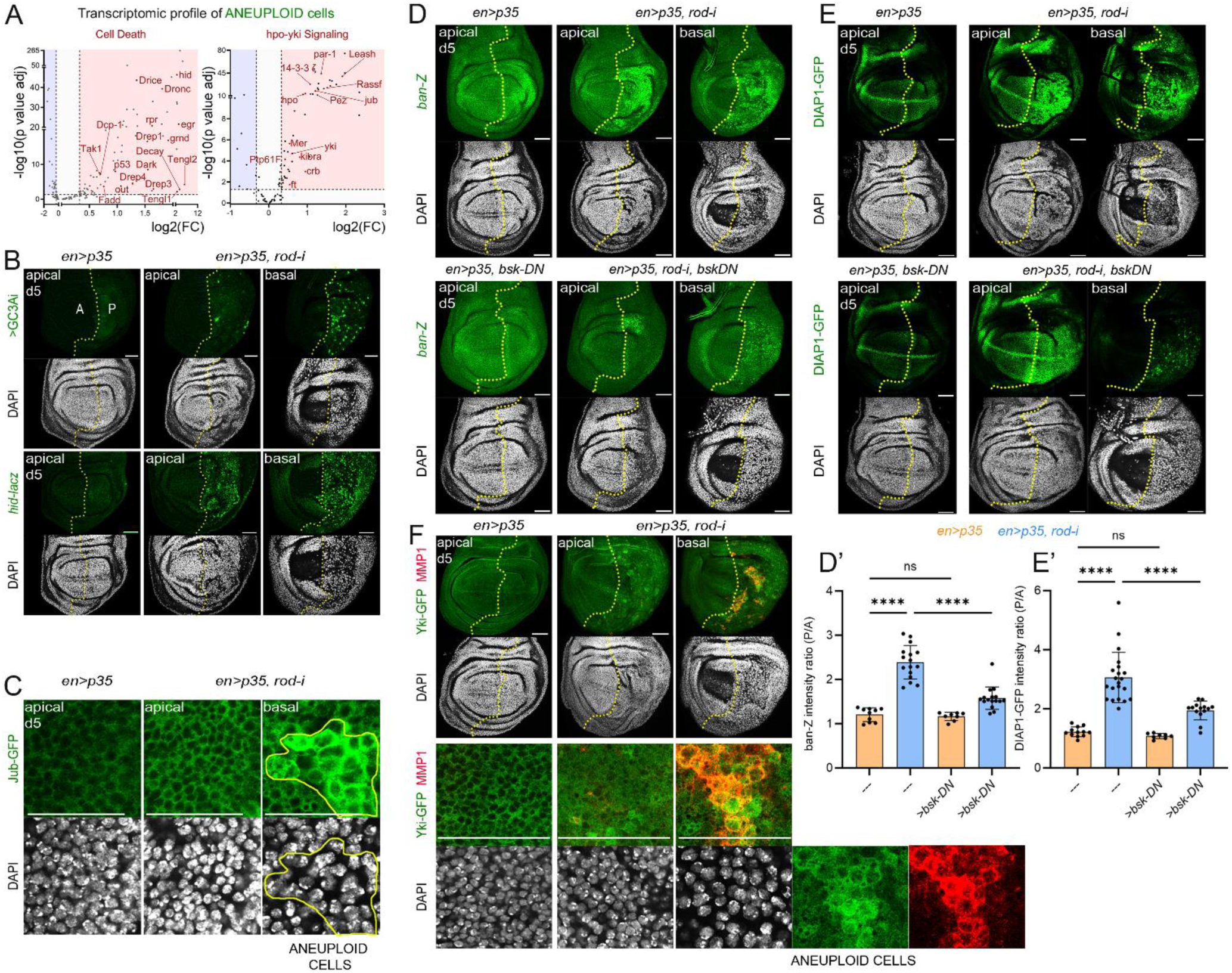
The transcriptome of aneuploidy-induced senescent cells points to activation of the apoptotic and Yorkie pathways. **(A)** Volcano plot representing selected genes in ‘Cell death - GO0008219’ (left), and ‘hpo-yki signaling - GO0035329’ (right) classes that are differentially upregulated (red area, FC>1.3 and p-value<0.05), downregulated (blue area, FC<-1.3 and p-value<0.05), or not differentially expressed (white area, p-value>0.05 or 1.3>FC>-1.3) in aneuploid cells when compared to P cells of control discs. (**B**-**F**) Control wing discs (first column) and wing discs subjected to CIN (remaining columns), stained to visualize GC3Ai or *hid-lacZ* (green, **B**), Jub-GFP (green, **C**), *bantam-lacZ* (green, **D**), DIAP1-GFP (green, **E**), Yorkie-GFP (green, **F**), DAPI (white, **B**-**F**), MMP1 (red, **F**), and expressing the indicated transgenes in the P compartment under the control of the *en-gal4* driver. In **C**, aneuploidy-induced senescent cells are marked by a yellow line. Scale bars, 50 µm. Discs were dissected 5 days AEL. The boundary between A and P cells is labelled by a yellow line. Note in **D** and **E,** a reduction in the expression levels of *bantam-lacZ* and *DIAP1-GFP* upon expression of a dominant negative form of JNK (Bsk-DN). (**D’**, **E’**) Histograms plotting the P/A signal intensity ratio of *bantam-lacZ* (**D’**) and *DIAP1-GFP* (**E’**) of wing discs expressing the indicated transgenes in the P compartment under the control of the *en-gal4* driver. Ordinary one-way ANOVA was performed to analyse the data. *p<0.05, **p<0.01, ***p<0.001, **** p<0.0001, ns, not significant. See also Table S3. See also Figure S3, and Tables S1 and S3.

**Figure 4.**
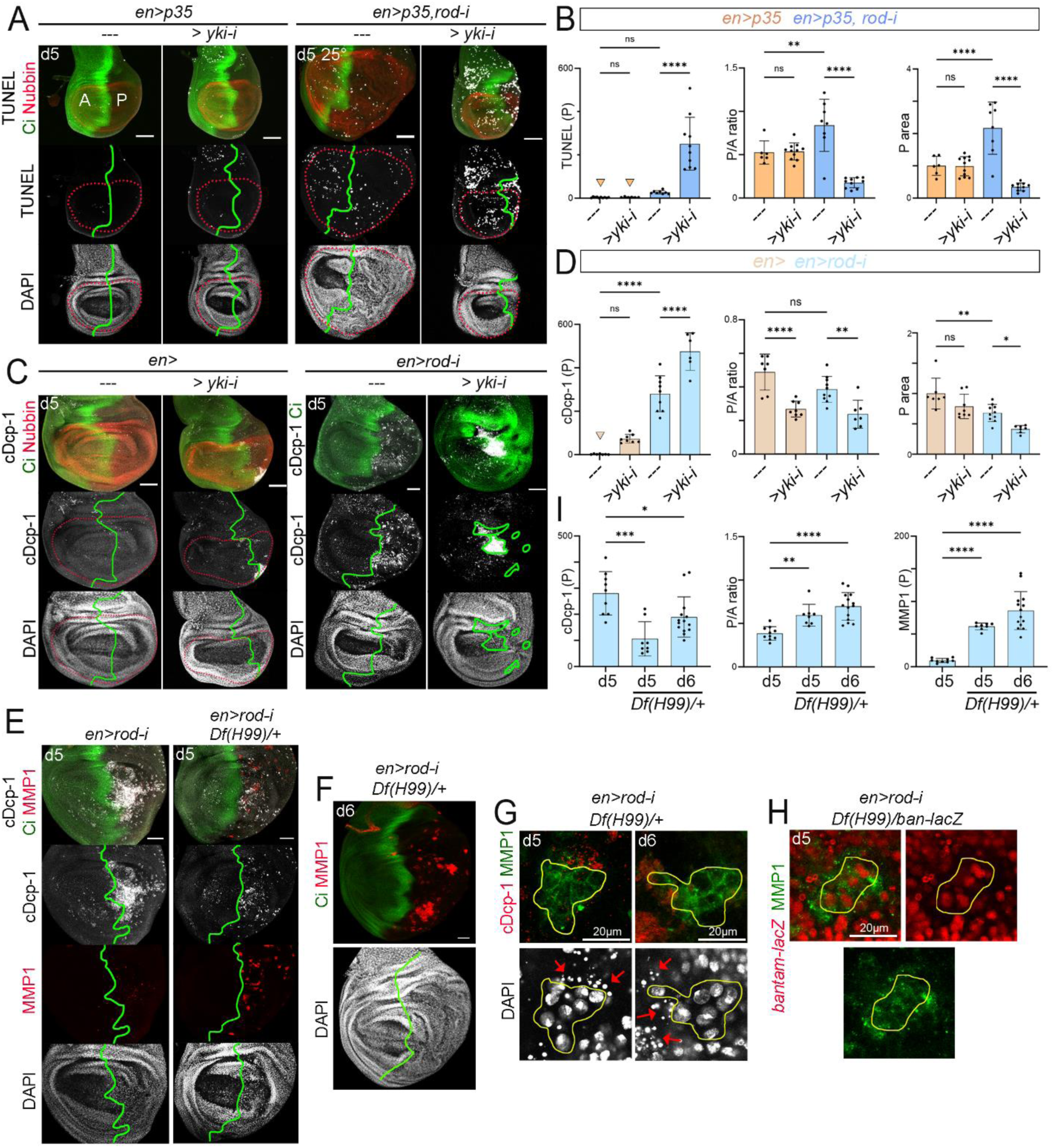
Yorkie plays a pro-survival role in CIN tissues. (**A, C, E, F, G, H**) Control wing discs (first columns in **A**, **C**) and wing discs subjected to CIN (remaining columns), stained to visualize TUNEL (**A**, white), cDcp-1 (**C**, **E**, white, **G**, red), *bantam-lacZ* (red, **H**), Ci (green, **A**, **C**, **E, F**), Nubbin (red, **A**, **C**), MMP1 (red, **E**, green, **G**, **H**), and DAPI (white, **A, C, E, F, G**) expressing the indicated transgenes in the P compartment under the control of the *en-gal4* driver. Scale bars, 50 µm, unless indicated otherwise. Discs were dissected at the indicated days of development. The boundary between A and P cells is labelled by a green line, and the wing pouch by a red dotted line. In **G**, **H**, high magnification images of the basal sections of CIN tissues, where aneuploidy-induced senescent cells are found and marked by a yellow line. Red arrows in **G** point to dying cells. (**B**, **D**, **I**) Histograms plotting the amount of TUNEL (**B**, left), MMP1 (**I,** right) and cDcp-1 signal (**D, and I,** left) in the P wing pouch compartment (normalized to the size of the corresponding compartment), the normalized size of the P wing pouch compartment (**B**, **D**, right), and size ratio between the P and A wing pouch compartments (**B, D and I,** centre) of wing discs expressing the indicated transgenes in the P compartment under the control of the *en-gal4* driver. Ordinary one-way ANOVA was performed to analyse the data plotted in **B, D, I;** * p<0.05, ** p<0.01, *** p<0.001, **** p<0.0001, ns, not significant. See also Table S3.

### A negative effect of CIN tumours on growth and survival of neighbouring cells

We noticed that the transcriptional profile of the neighbouring cell population (cells located in the nearby A compartment and not subject to CIN, Figure 1B) showed upregulation of genes involved in cell death and a reduction in translation activity (Figure S4A-C). We next examined the response of this cell population (A compartment) to the nearby hyperplastic tissue (P compartment) subject to CIN and additional blockage of the apoptotic pathway at different time points of development. As anticipated, the P compartment showed a steady increase in size, whereas the size of the A compartment was significantly reduced compared with controls (Figure 5A, quantified in Figure 5B). After eight days of development, the A compartment reached the expected size of a control A compartment, while the P compartment continued to grow. The differential growth dynamics of CIN and neighbouring cell populations were also observed when analysing the ratio of their corresponding sizes [P/A ratio; Figure 5B]. Interestingly, the negative impact of the CIN tissue on the growth dynamics of the A compartment was accompanied by an increase in the number of dying cells labelled by TUNEL staining (Figure 5A, quantified in Figure 5C), a reduction in mitotic activity labelled with pH3 antibody (Figure 5D, quantified in Figure 5C), and a decrease in EdU incorporation (Figure 5E, quantified in Figure 5C). Non-autonomous cell death was also observed upon induction of CIN in other regions of the wing primordium (Figure S4D).

**Figure 5.**
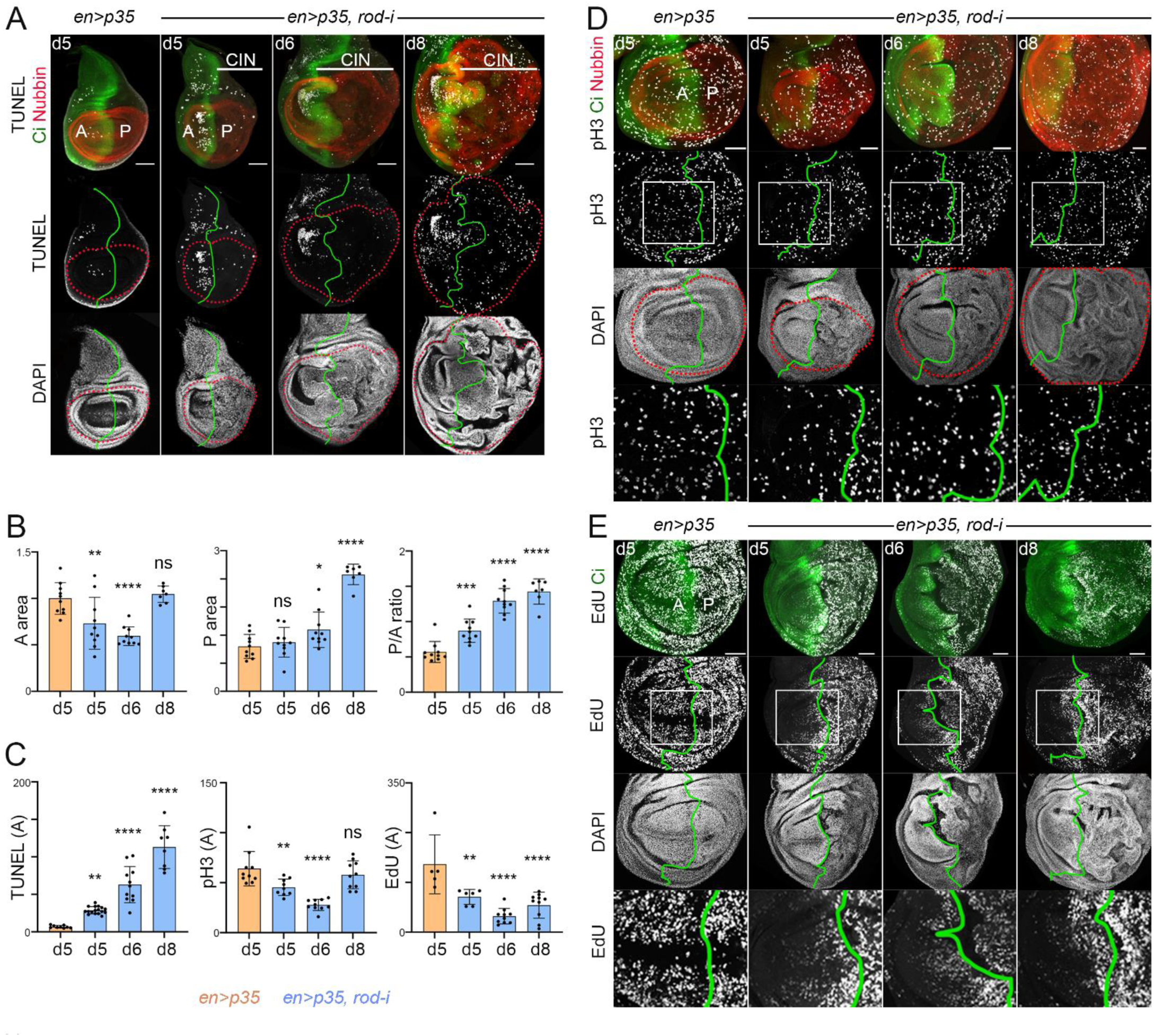
A non-autonomous effect of CIN tumours on the proliferation, growth, and survival of the neighbouring tissue. (**A**, **D**, **E**) Control wing discs (first column) and wing discs subjected to CIN (remaining columns), and expressing p35 in the P compartment under the control of the *en-gal4* driver, stained for Ci (green, to label the A compartment), DAPI (white), Nubbin (red, to label the wing pouch cells, **A**, **D**), TUNEL (white, **A**), pH3 (white, **D**), and EdU (white, **E**). Scale bars, 50 µm. Days of dissection after egg laying are as indicated. The boundary between A and P cells is labelled by a green line. The tissue subjected to CIN is marked in **A**. High magnification of the squared regions is shown in the lower panels in **D**, **E**. Note the non-autonomous induction of cell death (**A**) and block of cell proliferation (**D**, **E**) in A cells. (**B**, **C**) Histograms plotting in **B** the normalized size of the A and P compartments and the size ratio between these compartments, and in **C** the amount of TUNEL, pH3, and EdU incorporation in the A wing pouch compartment (normalized to the size of the corresponding compartment) of wing discs expressing the indicated transgenes in the P compartment under the control of the *en-gal4* driver. Ordinary one-way ANOVA was performed to analyse the data plotted in **B** and **C**. *p<0.05; **p<0.01; ***p<0.001; **** p<0.0001; ns, not significant. See also Figure S4 and Table S3.

We next addressed whether the observed response of the A compartment to the nearby CIN tissue was driven by any of the signalling molecules secreted by aneuploidy-induced senescent cells (Table 1). Interestingly, Dilp8, a molecule secreted by damaged tissues and well known to block the production of Ecdysone (*50*, *51*), a systemic hormone driving the larval to pupal transition and promoting proliferative growth, was among the most upregulated secreted proteins in aneuploidy-induced senescent cells (Figure 2A and Table 1). Using a GFP protein trap reporter, we confirmed the expression of Dilp8 in aneuploidy-induced senescent cells (Figure 6A). Depletion of *dilp8* in the CIN tissue (with a *UAS-RNAi*) restored mitotic activity and EdU incorporation levels, but not the number of apoptotic cells, in the adjacent A compartment (Figure 6B and C, quantified in Figure 6B’ and C’, Figure S5A, B). Interestingly, Dilp8 overexpression reduced mitotic activity and the size of the A compartment even further (Figure 6C, quantified in Figure 6C’). Since experimental manipulation of Dilp8 levels in the P compartment was unable to have a significant effect on the size of the nearby A compartment (Figure 6D), we searched for additional genes expressed in aneuploidy-induced senescent cells with a known role in inhibiting proliferative growth and found ImpL2, an orthologue of IGFBP7 that binds to and inhibits the function of *Drosophila* insulin-like peptides (*52*, *53*) and consequently the growth-promoting PI3K signalling pathway (Table 1), an upregulation that was confirmed with a GFP protein trap reporter (Figure 6A). Depletion of *ImpL2* in the CIN tissue (with a *UAS-RNAi*) restored mitotic activity and EdU incorporation levels (Figure 6B and C, quantified in Figure 6B’ and C’) in the adjacent A compartment but not number of apoptotic cells (Figure S5A, B), and ImpL2 overexpression reduced mitotic activity even further (Figure 6C, quantified in Figure 6C’). As observed in the case of Dilp8, experimental manipulation of ImpL2 levels had no significant impact on the size of the A compartment either (Figure 6D). Interestingly, co-depletion of both *dilp8* and *impL2* was able to dampen the reduction in the size of the A compartment caused by the nearby CIN tissue (Figure 6D). However, we noticed that the size of the P compartment was also dramatically increased upon *dilp8* and *impL2* co-depletion (Figure 6E), thus resulting in visibly bigger tumours (Figure 6F). These results are consistent with the previously reported systemic effects of these two secreted factors (*51*, *53*, *54*), and point to an additive tumour-suppressive role of these two systemic molecules, most probably by interfering with Ecdysone and dILPs, two growth-promoting systemic hormones. The reduction in the proliferative activity of the A compartment was not observed in those cells abutting the CIN tissue (Figure 5E and Figure S5D), thus pointing to a signal originating in the P compartment and counteracting the negative effects of Dilp8 and ImpL2 on proliferation. Indeed, the proliferative activity of these cells relies on the activity of the mitogenic molecule Wg produced by senescent cells (Table 1, (*29*), see also Figure S5C), as Wg depletion reduced the levels of EdU incorporation in these cells (Figure S5D, E).

**Figure 6.**
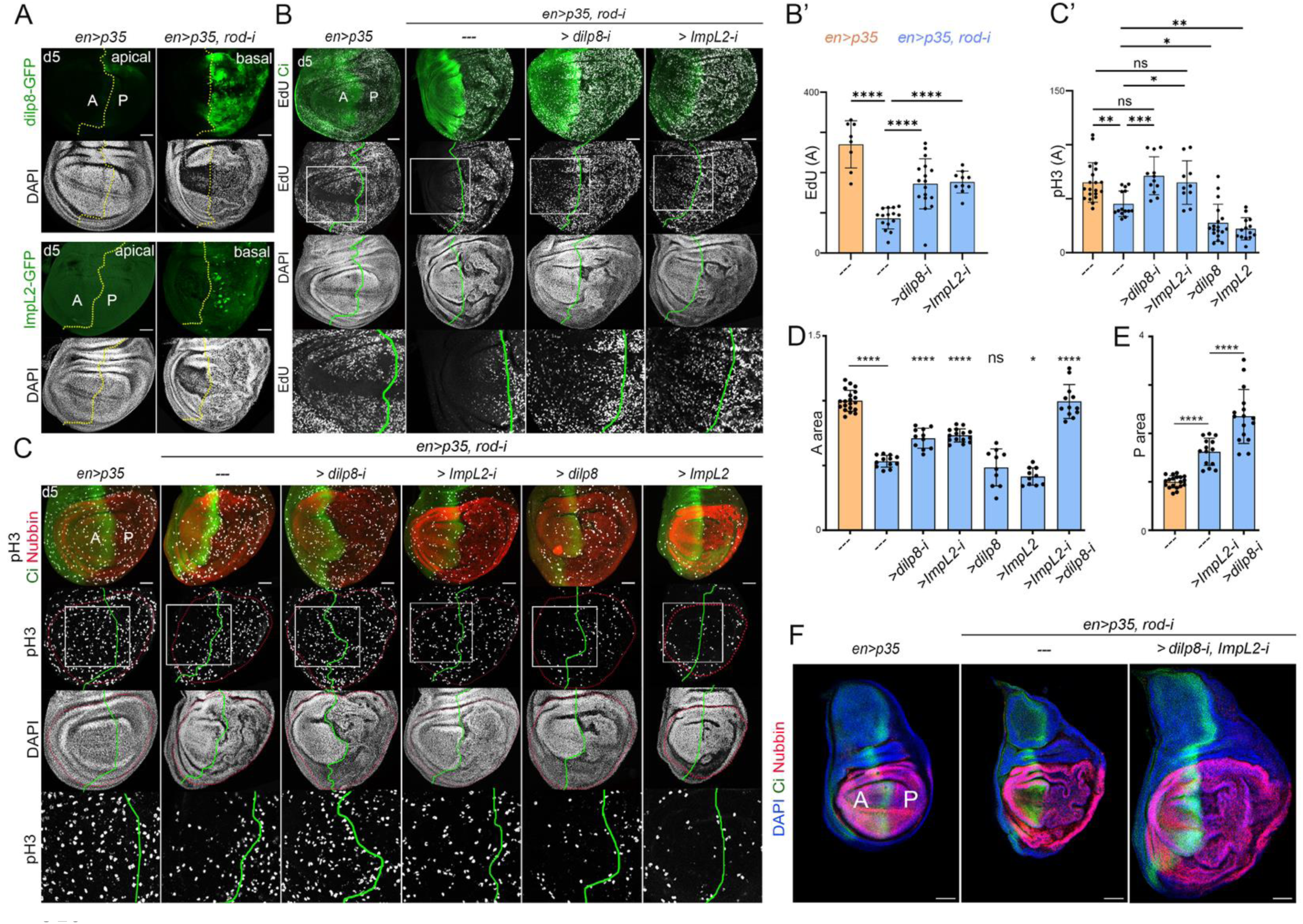
A growth suppressor role of Dilp8/Relaxin and ImpL2/IGFBP7 in CIN tumours. (**A-C, F**) Control wing discs (first column) and wing discs subjected to CIN (remaining columns), stained to visualize Dilp8-GFP (green, **A**), ImpL2-GFP (green, **A**), EdU (white, **B**), pH3 (white, **C**), DAPI (white in **A**-**C**, blue in **F**), Nubbin (red, to label the wing pouch cells, **C**, **F**), and Ci (green, to label the A compartment, **B**, **C**, **F**), and expressing the indicated transgenes in the P compartment under the control of the *en-gal4* driver. Scale bars, 50 µm. Discs were dissected 5 days after egg laying. The boundary between A and P cells is labelled by either a green line or a yellow dashed line. High magnifications of the squared regions are shown in the lower panels in **B**, **C**. (**B’, C’, D, E**) Histograms plotting the amount of EdU incorporation and pH3 in the A wing pouch compartment (normalized to the size of the corresponding compartment, **B’**, **C’**), and the normalized size of the A (**D**) and P (**E**) wing pouch compartments of wing discs expressing the indicated transgenes in the P compartment under the control of the *en-gal4* driver. Ordinary one-way ANOVA was performed to analyse the data plotted in **B’, C’, D, E.** * p<0.05; **p<0.01; ***p<0.001; ****p<0.0001; ns, not significant. See also Figure S5 and Table S3.

### A feed-forward loop driving tumour growth

The non-autonomous induction of cell death and reduction in tissue size caused by CIN tumours was, at least in part, a consequence of activation of the apoptotic pathway as halving the doses of the pro-apoptotic genes *reaper*, *hid* and *grim* (in *Df(H99)/+* larvae) caused a reduction in the levels of non-autonomous cell death (Figure 7D, D’) and restored the size of the nearby A compartment (Figure 7E). We noticed that targeted depletion of these three genes with the use of a synthetic micro-RNA [*UAS-miRHG*, (*55*)] only in the P compartment did not have any effect on the levels of cell death and on the size of the A compartment (Figure 7D, D’, E). In search of secreted molecules involved in the non-autonomous induction of the apoptotic pathway (Table 1), we noticed that aneuploidy-induced senescent cells showed upregulation, on the one hand, of elements of the JAK/STAT pathway, including ligands (cytokines Upd1, Upd2, and Upd3, Table 1) and the receptor Domeless (Dome, Figure 7A), and, on the other hand, the TNF-α ligand Eiger (Figure 2A and Table 1), which activates the JNK signaling pathway through its receptor Grindelwald (Grnd, (*56*)). Both JNK and JAK/STAT signalling pathways are known in CIN tissues to induce apoptosis through transcriptional induction of pro-apoptotic genes (*31*). We confirmed activation of the JAK/STAT pathway with a GFP transcriptional reporter (STAT-GFP (*57*)), and, most interestingly, we observed that it was also activated in the nearby A compartment (Figure 7B). We confirmed induction of *eiger* expression with a transcriptional reporter (Figure 7C, see also (*27*)). We next addressed the potential contribution of Upd ligands, the JAK/STAT pathway and Eiger to the effects of CIN tissues on the nearby wild type cell population.

**Figure 7.**
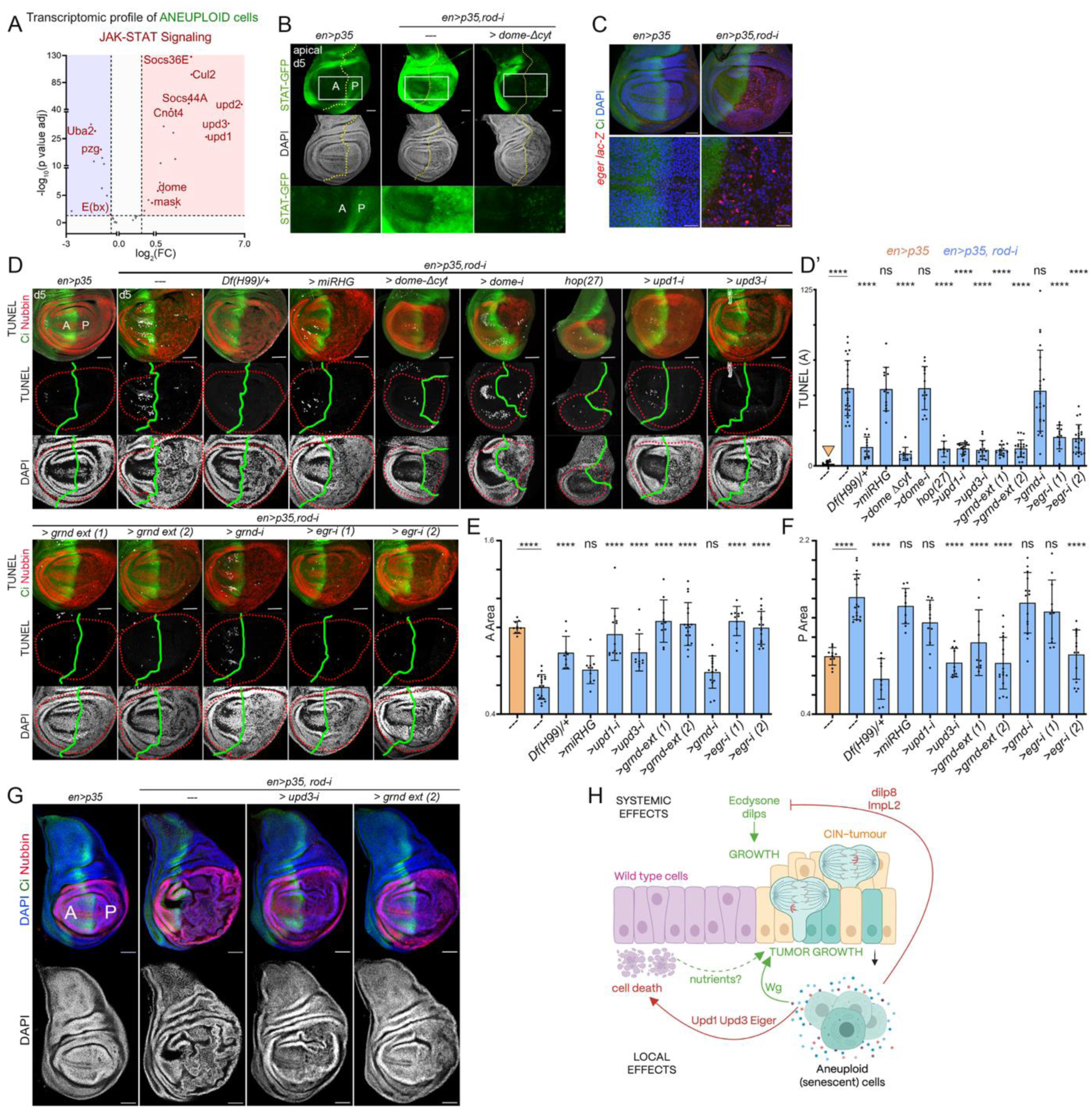
A feed-forward loop involved in the growth of CIN tumours. **(A)** Volcano plot representing selected genes in JAK-STAT Signaling - GO0097696 class that are differentially upregulated (red area, FC>1.3 and p-value<0.05), downregulated (blue area, FC<-1.3 and p-value<0.05), or not differentially expressed (white area, p-value>0.05 or 1.3>FC>-1.3) in aneuploid cells when compared to P cells of control discs. (**B, C, D, G**) Control wing discs (first column) and wing discs subjected to CIN (remaining columns) stained to visualize STAT-GFP (green, **B**), *eiger-lacZ* (red, **C**), TUNEL (white, **D**), Nubbin (red, to label the wing pouch cells, **D**, **G**), and Ci (green, to label the A compartment, **C, D, G**), and DAPI (white in **B**, **D**, blue in **C**, **G**) and expressing the indicated transgenes in the P compartment under the control of the *en-gal4* driver. Scale bars, 50 µm. Discs were dissected 5 days AEL. The boundary between A and P cells is labelled by a green or a yellow dashed line, or by Ci staining. High magnification of the squared regions are shown in the lower panels in **B**. (**D’, E, F**) Histograms plotting the amount of TUNEL (**D’**), and the size of the A (**E**) and P (**F**) wing pouch compartments of wing discs expressing the indicated transgenes in the P compartment under the control of the *en-gal4* driver. Ordinary one-way ANOVA was performed. *p<0.05, **p<0.01, ***p<0.001, **** p<0.0001, ns, not significant. (H) Model: in epithelial tissues subjected to restricted induction of CIN in the P cell population (yellow), aneuploid cells (green) delaminate basally and enter a senescent state. Members of the SASP impact on the neighbouring wild type cell population (the A cell population depicted in purple) locally of systemically to induce cell death and regulate cell proliferation and growth. Non-autonomous cell death feeds back to the tumour to enhance its growth. Cartoon was generated with Biorender.com. See also Tables S1 and S3.

Expression of a truncated form of receptor Domeless lacking the intracellular domain in the CIN tissue (Dome-ΔCYT) led to a marked reduction in the levels of STAT-GFP signalling, not only cell autonomously but also in the nearby A compartment (Figure 7B). This finding is consistent with the proposed role of Dome-ΔCYT in blocking JAK/STAT signalling by trapping the Upd ligands and competing with the endogenous receptor (*30*). Interestingly, the non-autonomous reduction in STAT-GFP levels caused by Dome-ΔCYT was accompanied by a decrease in the number of apoptotic cells (Figure 7D, quantified in Figure 7D’). In contrast, depletion of Dome by RNAi, which is expected to block ligand-dependent JAK/STAT signalling in a cell autonomous manner without trapping the Upd ligands, did not affect the neighbouring A compartment (Figure 7D, quantified in Figure 7D’). We found that both transgenes (*UAS-dome-ΔCYT* and *UAS-dome-RNAi*) caused a similar reduction in the size of the P compartment, most probably caused by the developmental role of the JAK/STAT pathway in regulating the growth of this compartment (*58*). The non-autonomous reduction in the number of apoptotic cells caused by Dome-ΔCYT was also observed in tissues mutant for *hop* (the *Drosophila* JAK orthologue) and upon depletion of the cytokines *upd1 or upd3* by RNAi in CIN and aneuploid cells (Figure 7D, quantified in Figure 7D’). We observed that neither *upd1* depletion nor Dome-ΔCYT expression in CIN tissue ameliorated the non-autonomous reduction in proliferation dynamics (Figure S6A, quantified in Figure S6B).

In a similar manner, depletion of *eiger* or expression of a dominant negative version of its receptor Grnd (Grnd-extra) in CIN tissues, known to trap Eiger ligand thus blocking its function (*56*), reduced the number of apoptotic cells in the nearby A compartment (Figure 7D, quantified in Figure 7D’). Depletion of *grnd* by RNAi, which is expected to block Eiger-dependent JNK activation in a cell autonomous manner without trapping the Eiger ligand, did not affect the apoptotic levels in the neighbouring A compartment (Figure 7D, quantified in Figure 7D’). Most interestingly, depletion of Upd1, Upd3 or Eiger rescued the reduction in the size of the A compartment (Figure 7E). Similar results were observed with Grnd-extra (Figure 7E). All these results point to an additive role of multiple molecules secreted from aneuploidy-induced senescent cells and acting in nearby wild type tissues to induce cell death and compromise growth.

We next analysed whether the non-autonomous cell death had any impact on the growth of the tumour itself. Interestingly, the reduction in the levels of non-autonomous cell death and the restoration of the size of the A compartment caused by depletion of Eiger, the Upd1 and Upd3 cytokines or expression of their truncated receptors was in all cases accompanied by a reduction in the size of the hyperplastic tissue (P compartment) subject to CIN and additional blockage of the apoptotic pathway (Figure 7F, G). We noticed that depletion of *grindelwald* by RNAi did not have any impact on the size of the P compartment, thus ruling out an autocrine role of Eiger in CIN tumors [Figure 7F, see also (*27*)]. Depletion of several genes of the JAK/STAT signalling pathway by RNAi was previously shown to have no impact on the size of CIN tumours either, thus ruling out also an autocrine role of Upd cytokines (*31*). These results indicate that several SASP members act in an additive manner on neighbouring tissues to reduce growth through cell death and that these non-autonomous effects feed back to the tumour to reinforce its growth.

Although the classical apoptotic pathway appears to play a major role in the non-autonomous induction of cell death and reduction in tissue size exerted by Upd cytokines and Eiger, as cell death and tissue size were both largely rescued by halving the doses of the pro-apoptotic genes (Figure 7D, D’, E), non-apoptotic cell death might also contribute to the observed effects on tissue size. On the one hand, Eiger and the JNK pathway have been reported to induce also non-apoptotic cell death through the production of ROS (*59*). On the other hand, Upd cytokines and the JAK/STAT pathway, well known to induce autophagy in CIN tissues (*19*), might also cause cell death through autophagy. Consistent with this proposal, we noticed that in those regions within the nearby A compartment with the highest levels of apoptotic activity (monitored with an antibody to detect the cleaved version of the effector caspase Dcp1, cDcp1), autophagy was also induced, as reflected by the accumulation in puncta of a mCherry-tagged Atg8a under the control of the endogenous Atg8a promoter (Atg8a; Figure S7A). This induction was largely dependent on Upd1 derived from CIN tissue, as *upd1* depletion led to a significant reduction in the amount of Atg8a signal in the A compartment (Figure S7A, A’). Compromising autophagic flux, by supplementing the larval food with chloroquine, led to a significant reduction in the extent of cell death observed in the A compartment (Figure S7B, B’). We noticed that caspase activation in CIN cells, which play a non-apoptotic role by inducing cell motility in aneuploid cells, was not reduced with this treatment, which is consistent with their reported activation through transcriptional induction of pro-apoptotic genes by JNK and JAK/STAT signalling pathways (*31*). Overall, these observations point to the potential contribution of different forms of cell death in the observed feed-forward loop involved in the growth of CIN tumours.

## Discussion

### The transcriptome of aneuploidy-induced senescence

From flies to mammals, CIN is widely recognized as a driver of cellular senescence, primarily through the generation of highly aneuploid karyotypes (*19–22*), and cellular senescence and its associated senescence-associated secretory phenotype (SASP) are major triggers of inflammation and tumorigenesis (*60*). Here we have characterized the transcriptional profile of aneuploidy-induced senescence in an epithelial model of CIN in *Drosophila* that recapitulates most emerging cellular behaviours observed in mammalian epithelial tissues upon CIN (*10*, *22*, *24–28*, *61–63*). Thus, in fly epithelial tissues subjected to CIN and upon additional blockade of the apoptotic pathway, aneuploid cells are extruded from the epithelium and enter a state of senescence characterized by cell cycle arrest in G2, increased motility, and a highly secretory phenotype that drives the hyperplastic growth of the epithelium in a non-autonomous manner (*64*). Consequently, CIN tissues acquire a tumour-like, unlimited growth potential and highly invasive behaviour. All these processes rely on the activity of the JNK signalling pathway. As it occurs from yeast to mammals, aneuploidy activates autophagy as the primary quality control mechanism for maintaining proteostasis in response to gene dosage imbalance and proteotoxic stress (*65*, *66*). Saturated use of the autophagy machinery in these cells leads to the accumulation of dysfunctional mitochondria as a result of compromised mitophagy, and these mitochondria become active sources of ROS (*19*), which ultimately activate JNK via Ask1 (*27*). The use of the GAL4/UAS system to trigger CIN in only one half of the growing wing epithelium and a GFP transcriptional reporter of JNK (MMP1-GFP, (*39*)) to track aneuploid cells has allowed us to characterize the transcriptional profile of aneuploidy-induced senescent cells, the epithelial layer subjected to CIN that enters a hyperplastic mode of growth, and the nearby wild-type population. Remarkably, most cellular responses to aneuploidy and senescence are robustly reflected at the transcriptional level, as shown by the up- or down-regulation of a large number of genes involved in any of these responses (Figures 1 and 2). Thus, cell cycle progression is locked in aneuploid cells due to the downregulation of most genes involved in DNA replication (including all subunits of the DNA polymerase, and Orc and Mcm complexes) and G2/M transition (including *Dcdc25*/*stg*, *cdk1*, *cycA*, and *cycB*). The induction of autophagy, a process that delivers its cargo to lysosomes for degradation, is accompanied by the upregulation of many autophagy (*atg4a*, *atg8a*, *atg9*, *atg17,* and *atg18b*) and lysosomal genes (e.g., *Lamp1*, *Vps13D*, *Vps16A, Vamp7,* and *lqf*). Similarly, the reported ability of aneuploid cells to become motile through modulation of the actin cytoskeleton is accompanied by the upregulation of many actin cytoskeleton proteins, including Sqh, Cortactin, Rok, Fak, and Arpc2-5. One of the most remarkable properties of senescent cells is the SASP (*67*). Our transcriptomic analysis has unravelled a significant upregulation of genes involved in secretion (e.g., *Tango 5, 11, Sec 5, 6, 33, 24CD,* and *Rab 1, 2, 6, 7, 10,* and *35*), reinforced with the use of cellular reporters to unveil an enlarged secretory apparatus, and has allowed us to identify all the elements of the SASP, where 10% of the most upregulated genes in these cells encode for secreted proteins (Table S2). Remarkably, many of the cellular behaviors observed in our fly model of CIN have been observed in recent years in mammalian tissues, thus reinforcing its translational impact. Thus, mammalian cells with complex karyotypes caused by chromosome segregation defects are also extruded from the epithelium (*61*), and they are cell cycle-arrested, exhibit features of senescence, and produce pro-inflammatory signals (*22*). Most interestingly, CIN also drives tumorigenesis in mammals [e.g. (*62*)] and promotes metastasis (*10*).

### An anti-apoptotic role of Yorkie in senescent cells

*Drosophila* CIN tissues are exposed to the activation of pro-apoptotic pathways JNK and JAK/STAT that lead to the elimination of aneuploid cells in the tissue (*24*, *31*). It is the blockage of the apoptotic pathway by the expression of the baculovirus protein p35 what unravels a subversive role of JNK in driving cellular senescence, and JAK/STAT in promoting cell motility. However, the apoptotic pathway is not completely blocked under these conditions as shown by the high activity levels of effector caspases in the tissue, which have been shown to contribute in a non-apoptotic manner to the invasive behaviour of aneuploid cells. These results point to the presence of an endogenous pathway that counteracts the pro-apoptotic activities of JNK and JAK/STAT. Similar to how senescent cells in mammals resist normal apoptotic pathways through expression of the Bcl2 anti-apoptotic protein (*68*), our results indicate that the Hippo/Yorkie pathway plays an analogous role in senescent fly cells. Activation of this pathway is most probably the result of JNK activation and cellular tension positively regulating the cortical localization of Ajuba expression to sequester Warts, thus allowing Yorkie to translocate into the nucleus and drive gene expression (*47*, *69*). Although DIAP1 (a negative regulator of the initiator Caspase Dronc (*43*)) or bantam miRNA (which acts negatively on the hid 3’UTR (*70*)) could be two potential target genes of Yorkie that block the death of senescent cells, the finding that effector caspases show high activity in these cells suggests that Yorkie negatively impinges on the apoptotic pathway even at the level of cell death execution. These experimental data contribute to the identification of Yorkie as a potential therapeutic strategy to block CIN-induced tumorigenesis by targeting senescent cells.

### The SASP mediates a tumour-host feed-forward loop: a case of super-competition

In our fly epithelial model of CIN, senescent cells show a remarkable ability to act at a paracrine, autocrine, and systemic level to drive tumour growth, invasiveness, and malignancy, respectively ^2^. They exert their action through elements of the SASP, including mitogenic molecules Wg and Wnt6, Matrix metalloproteinase MMP1, and cytokines such as Upd3. Here we identify a new axis of interaction between senescent cells with the nearby wild type cell population that contribute to tumour growth (Figure 7H). While the CIN tissue enters a hyperplastic mode of growth in response to the mitogenic molecules Wg and Wnt6 secreted by senescent cells, the nearby wild type cell population shows a reduction in growth rates, which is accompanied by a reduction in proliferative activity and an increase in the number of apoptotic cells. We identified several components of the SASP driving the observed non-autonomous effects: the Relaxin-like molecule Dilp8, the Insulin-like growth factor antagonist ImpL2 (the fly ortholog of IGFBP-7), the Upd1 and Upd3 cytokine and the TNF-α-orthologue Egr. While Dilp8 and ImpL2 were shown to be required and sufficient to drive the non-autonomous reduction in proliferative activity, Upd1, Upd3 and Eiger were found to be responsible for the reduction in growth and increase in apoptotic levels of the neighbouring cell population. We present evidence that Upd1 and Upd3 exerts its action by triggering the JAK/STAT signalling pathway and by inducing cell death. Consistent with the reported activities of Dilp8 and ImpL2 as systemic signals blocking the production or activity of those systemic hormones required for organismal growth [eg. ecdysone and dILPs, (*34*, *50*, *51*, *71*)], their negative impact on the proliferative activity was found to be extended to the tumour itself and not just to neighbouring tissues. These data expand the repertoire of signalling activities exerted by senescent cells and provide another example of non-cell-autonomous induction of cell death in the tumour host (*72*). Most interestingly, we present evidence that the non-cell-autonomous induction of cell death induced by Upd cytokines and Eiger is required to feed back to the tumour to enhance its growth, either as a consequence of the liberation of nutrients and metabolites, or the expression of signalling molecules by dying cells before being captured by circulating macrophages (Figure 7H). These tumour-host interactions parallel interface interactions observed during super-competition, an aggressive form of cell competition where a cell acquiring a specific mutation (often oncogenic) becomes “super-fit” and actively eliminates neighbouring, healthy wild-type cells to expand its territory (*35–37*).

## Materials and Methods

### Fly maintenance, husbandry and transgene expression

Strains of *Drosophila melanogaster* were maintained on standard medium (4% glucose, 55 g/L yeast, 0.65% agar, 28 g/L wheat flour, 4 ml/L propionic acid, and 1.1 g/L nipagin) at 25°C in light/dark cycles of 12 h. The resulting larvae after the egg laying (AEL) at 25°C were either transferred to 29°C, or maintained at 25°C for specific experiments as mentioned in the figures, and were dissected on the day AEL as mentioned in the figures. The sex of experimental larvae was not considered relevant to this study and was not determined. The strains used are summarized in Table S4.

### Standard induction of CIN, and endogenous labelling of different tissue subpopulations for transcriptomic analysis

Virgin female flies carrying *UAS-myrT, MMP1-GFP* transgenes were crossed with males carrying either *en-GAL4, UAS-p35* (Control) or *en-GAL4, UAS-p35, UAS-rod^RNAi^* (CIN) transgenes. Fertilized females were allowed to lay eggs at 25^°^C, and the resulting larvae were switched to 29^°^C 24 h AEL. Wing imaginal discs were then dissected after either 5 days (in case of control) or 7 days (in case of CIN) of induction at 29°C. Wing imaginal discs (40–50 in case of controls, and 30–40 in case of CIN) were cleaned in ice-cold 1X PBS and were incubated in 10X trypsin-EDTA (Sigma-Aldrich) at room temperature (RT) for 1 h, with regular nutation, and gentle pipetting. The single cell suspension was then subjected to centrifugation at 2500*g* for 2.5 min. The supernatant was gently removed, and the cells were stained with 1 ml of 1ug/ml of DAPI solution for 10 min at 4°C in low-light conditions, after resuspending the cell pellet. The cells were immediately transferred to flow-tubes, and FACS sorting was carried out on the different tissue subpopulations.

### FACS sorting of different tissue subpopulations

Flow cytometry experiments were carried out using a FACS Aria Fusion sorter (Beckton Dickinson, San Jose, California) equipped with a 4-laser configuration (405-488-561-640nm). Single cells were gated according to scatter parameters, using FSC width to avoid doublets. Dead cells were excluded using DAPI as a viability marker; only live single cells were analysed, and these were sorted on the basis of their fluorescence pattern. For sorting different CIN subpopulations from CIN tissues, P(Aneuploid) cells were distinguished as GFP+ RFP+ cells and P(CIN) cells as GFP-RFP+ cells, and the remaining GFP-RFP-cells were labelled as A(Aneuploid) cells. Similarly, for sorting the two subpopulations from the control tissues, RFP+ cells were attributed as P(control), while the RFP-cells were attributed as A(control). Fluorescence of GFP was detected using blue (488 nm) laser excitation and collecting green fluorescence using a 530/30 BP filter. RFP fluorescence was obtained using green-yellow laser excitation (561 nm) and collecting fluorescence using a 610/20 BP filter. A dot plot combining GFP and RFP emission on single live cells was used to select them for sorting into 1.5 ml Eppendorf tubes. A 70 µm nozzle was used to minimize the sorted volume into collection vials. During the entire experiment, the cells were under 300 RPM agitation and were maintained at 4°C. For each subpopulation, 30000 cells were collected in RNase-free 1.5 ml microcentrifuge tubes containing 10 µl of freshly prepared 2X Lysis Buffer following the methods from (*73*).

### RNA purification, library preparation, and bulk RNA-sequencing

Total RNA from a total of 4 replicates of 5 distinct subpopulations was then purified at the IRB Barcelona Functional Genomics Core Facility, as described in (*73*). mRNA was isolated from total RNA using the NEBNext Poly(A) mRNA Magnetic Isolation Module Kit (New England Biolabs). NGS libraries were prepared from the purified mRNA using the NEBNext Ultra II RNA Library Prep Kit for Illumina (New England Biolabs). A total of 18 cycles of PCR amplification were applied to all libraries. The final libraries were quantified using the Qubit dsDNA HS assay (Invitrogen) and quality controlled with the Bioanalyzer 2100 DNA HS assay (Agilent). An equimolar pool was prepared with the 20 libraries and submitted for sequencing at the *Centro Nacional de Análisis Genómico* (CNAG). A final quality control by qPCR was performed by the sequencing provider before paired-end 150 nt sequencing on a NovaSeq6000 S4 (Illumina). More than 275 Gbp of reads were produced, with a minimum of 39 million paired-end reads per sample.

### Data processing

Adapters from reads were removed using trim galore (v.0.6.7) (https://github.com/FelixKrueger/TrimGalore) with default parameters. Resulting reads were aligned to the dm6 *D. melanogaster* genome version using STAR (v.2.7.10a) (*74*). The count matrix was generated in R (https://www.R-project.org/) through the function “featureCounts” of the Rsubread package (*75*).

### Differential expression

Differential expression was performed using the DESeq2 package (*76*). For plotting purposes and generating principal components, the count matrix was normalized using the “vst” function.

### Gene set enrichment analysis

Gene set enrichment analysis was performed using the roastgsa package (*77*). GO and KEGG pathways were downloaded using the org.Dm.eg.db package.

### Immunohistochemistry

Larvae were selected according to the day AEL mentioned in the individual figures, and wing imaginal discs were dissected in phosphate-buffered saline (PBS), fixed for 20 min in 4% formaldehyde in PBS, and stained with antibodies in PBS with 0.3% BSA, 0.2% Triton X-100. The antibodies and reagents used in this study are summarized in Table S4.

### DNA synthesis

The Click-iT™ Plus EdU Alexa Fluor™ 647 Imaging Kit from Invitrogen (C10640) was used to measure DNA synthesis (S phase) in proliferating cells, following the manufacturer’s instructions. EdU (5-ethynyl-2’-deoxyuridine) provided in the kit is a nucleoside analogue of thymidine and is incorporated into DNA during active DNA synthesis. Time of incubation with EdU: 15 min

### TUNEL

Larvae were dissected in cold 1X PBS and fixed in 4% FA for 20 min. After fixation, three fast washes with PBT 0.3% Triton (PBS 1%, Triton X-100 0.3%) followed by four 15-min washes with PBT 0.3% Triton were performed. The samples were blocked with BBT (0.3% BSA, 0.3% Triton X-100) for 1 h rotating at RT and then incubated overnight with the primary antibody at 4℃. The next day, samples were subjected to three fast washes with PBT 0.3% Triton, and then they were washed four more times with PBT 0.3% Triton for 20 min each, followed by incubation with secondary antibodies diluted in BBT for 90 min rotating in the dark at RT. After incubation, the samples were subjected to three rounds of fast washes with PBT 0.3% Triton and four 20-min washes with PBT 0.3%. Samples were kept in BBT blocking overnight at 4°C. The following day, samples were permeabilized with 495 μl of 100 mM Na-Citrate and 2.5 μl of 20% TritonX for 30 min at 65°C and then subjected to three fast washes with PBT 0.3% Triton. They were then incubated twice for 5 min at RT with 50 μl of Equilibration buffer. Then 51 μl of TUNEL reaction mix (50 μl of TUNEL reaction buffer Component A, 1 μl TdT Enzyme, Biotium) was added and the samples were incubated for 2 h at RT. After incubation, the samples were subjected to three fast washes with PBT 0.3% Triton, followed by a 30-min incubation with DAPI at RT. Finally, three fast washes and three 10-min washes with PBT 0.3% Triton were performed. Stained tissues were kept in mounting media.

### Confocal microscopy

Samples were analysed using the following confocal microscopes: STELLARIS Confocal Platform with White Light Laser (Leica Microsystems), LSM 780 Zeiss, and LSM880 Zeiss fitted with a Fast Airyscan module (Carl Zeiss). Z-stacks were acquired using either the laser scanning confocal mode or the High-Resolution mode (Airyscan). Image acquisition was done with 20x, 25x, 40x, 63x, and 100x lenses. The most representative image is shown in all experiments.

### Chloroquine treatment

Flies of desired genotypes were crossed and set up at 25°C and shifted to 29°C 8 h AEL. After 48 h at 29°C, a working concentration of 3 mg/ml of chloroquine (with water as controls) was added to the food. Larvae were processed 72 h thereafter. Experimental flies and control individuals were grown in parallel.

### Tissue size, cell death and mitotic activity

The sizes of the whole wing disc (based on DAPI staining), the anterior (A) and posterior (P) compartments (based on Ci expression, in Figures 3-5) and the anterior (A) and posterior (P) areas of the wing pouch (labelled by Nubbin expression, in Figure 6 and 7) were measured manually using Fiji Software (NIH, USA) on apical sections (epithelium) of the wing disc obtained using a Zeiss LSM780 confocal microscope. A and P areas were normalized to the size of the respective control compartments. The left column of each graph represents the genotype which was considered as the control. The area of the signal to be quantified (TUNEL, cDcp-1, EdU, pH3, and mCh-Atg8) was set by manually adjusting the threshold and normalized to the area of the indicated larval territory (A or P compartments within the wing pouch) after making a maximum intensity Z-Projection.

### Statistical analysis

Statistical analysis was performed by either One-way ANOVA when comparing more than two experimental groups or an unpaired equal-variance two-tailed Student’s t-test when comparing the difference in means of a single experimental group with a control. Differences were considered significant when p values were less than 0.0001 (****), 0.001 (***), 0.01 (**), or 0.05 (*). All genotypes included in each histogram or scatter plot were subjected to the same experimental conditions (temperature and time of transgene induction) and analysed in parallel. Mean values and standard deviations were calculated, and the corresponding statistical analysis and graphical representations were carried out with GraphPad Prism 10.0 statistical software.

## Data availability

Materials generated for the study are available from the corresponding author on request. Source data are provided with this paper. The raw and processed RNAseq data have been deposited in the GEO database under accession number GSE306996.

## Acknowledgments

We thank E. Bach, J. Castelli-Gair, N. Tapon, M. Suzanne, G. Juhasz, the Bloomington *Drosophila* Stock Center (USA), the Vienna *Drosophila* Resource Center (Austria), and the Developmental Studies Hybridoma Bank (USA) for flies and antibodies, and IRB Barcelona’s Advanced Digital Microscopy, Functional Genomics, and Biostatistics and Bioinformatics Facilities for help. This work was funded by the PID2019-110082GB-I00 and PID2022-137673NB-I00 grants from the Spanish Ministry of Science, Innovation and Universities (MICIU) and the ERDF “Una manera de hacer Europa”, and the European Union’s Horizon 2020 research and innovation programme under the Marie Skłodowska-Curie grant agreement number 945352. We gratefully acknowledge institutional funding from the MICIU through the Centres of Excellence Severo Ochoa Award, and from the CERCA Programme of the Catalan Government.

## Author Contributions

All the authors conceived and designed the experiments and analysed the data. K.G. and A.K. performed the experiments, M.M. and K.G. prepared the final figures, and M. M. supervised the whole project and wrote the paper.

