## Supplementary material for "A tumour-host feed-forward loop contributes to malignant growth in chromosomal instability-induced epithelial tumours": ms & figures & Table 1

### Supplementary Figures and Tables

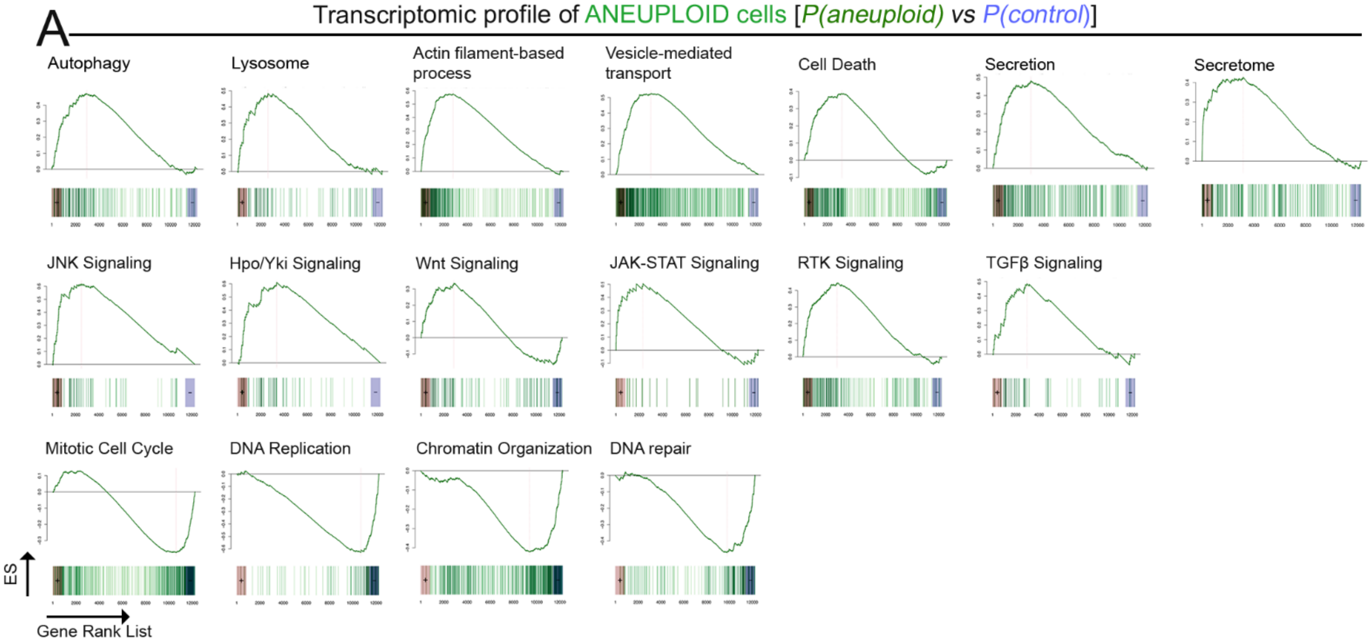


**Figure S1. Transcriptomic landscape of aneuploidy-induced senescent cells (**related to Figure 1**)**

**(A)** Mountain plots representing Enrichment Score (ES) upon Gene Set Enrichment Analysis (GSEA) comparing P(Aneuploid) with P(control) cells. Top row - up regulated cellular processes: Autophagy - GO0006914, Lysosome - GO0005764, Actin filament-based process - GO0030029, Vesicle-mediated transport - GO0016192, Cell death - GO0008219, Secretion - GO0046903, Secretome - Custom Gene Set. Middle row - up regulated signaling pathways: JNK Signaling - GO0007254, Hpo/Yki Signaling - GO0035329, Wnt Signaling - GO0198738, JAK-STAT Signaling - GO0097696, RTK Signaling - GO0007169, TGFβ Signaling - GO0030509. Bottom row - down regulated cellular processes: Mitotic cell cycle - GO0000278, DNA Replication - GO0006260, Chromatin Organization - GO0006325, DNA repair - GO0006281. See also Table S1.

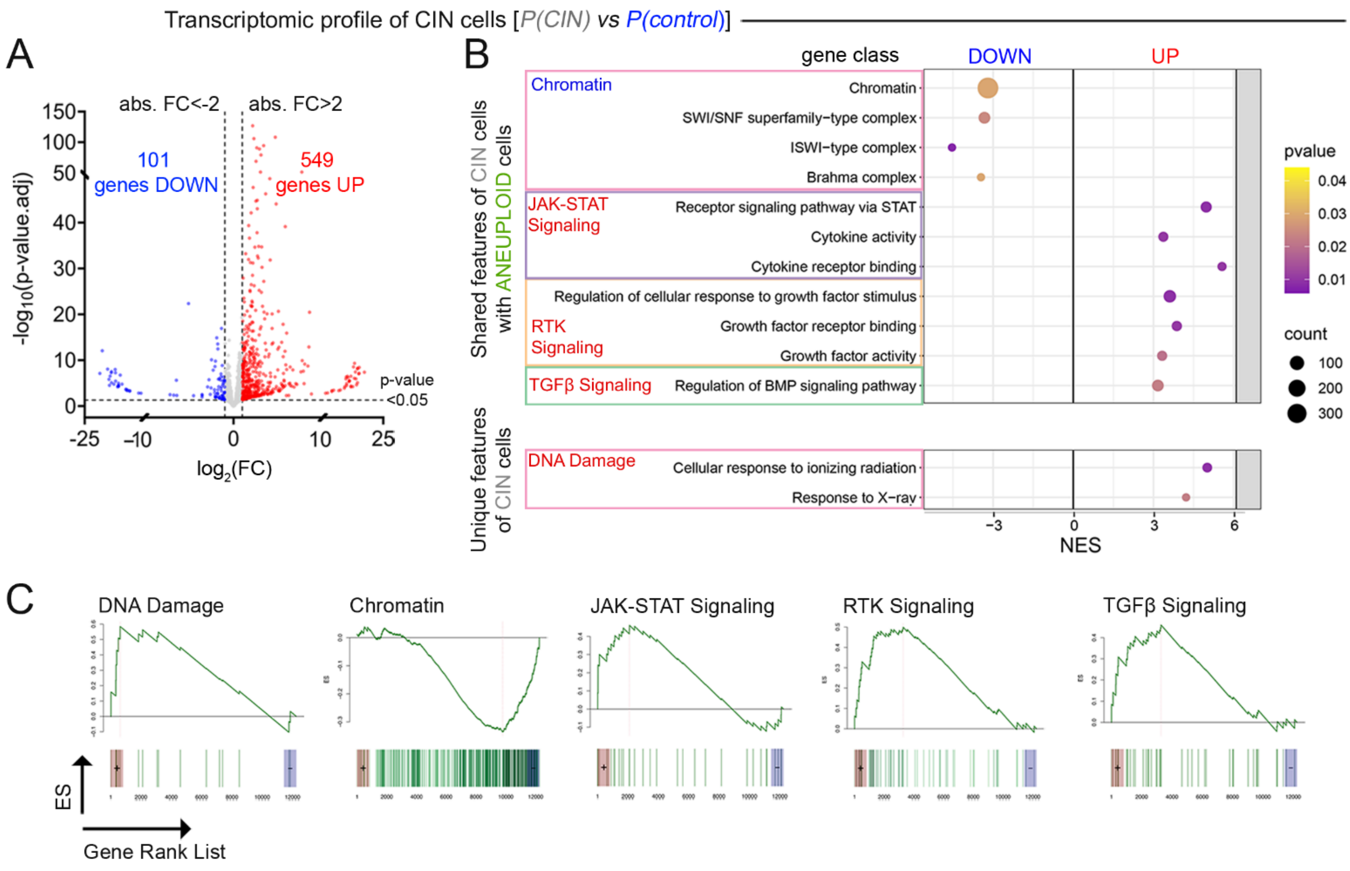


**Figure S2. Transcriptomic landscape of proliferating epithelium subjected to CIN (**related to Figure 1**)**

**(A)** Volcano plot representing all genes that are differentially up- (red dots, FC>2 and p-value<0.05) or down-regulated (blue dots, FC<-2 and p-value<0.05), or not differentially expressed (grey dots, p-value>0.05 or 2>FC>-2) in proliferating cells subjected to CIN when compared to P cells of control discs. **(B)** Bubble plot representing Gene Ontology (GO) enrichment analyses of proliferating epithelium subjected to CIN when compared to P cells of control discs. Size of the bubble represents the number of measured genes and color-scale represents the associated p-value. On the top, GOs shared with aneuploid cells. On the bottom, those GOs uniquely upregulated in the proliferating epithelium subjected to CIN. **(C)** Mountain plots representing Enrichment Score (ES) upon Gene Set Enrichment Analysis (GSEA) of proliferating cells subjected to CIN when compared to P cells of control discs. GO classes: DNA Damage - GO0071479, Chromatin - GO0000785, JAK-STAT Signaling - GO0097696, RTK Signaling - GO0090287, TGFβ Signaling - GO0030510. See also Table S1.

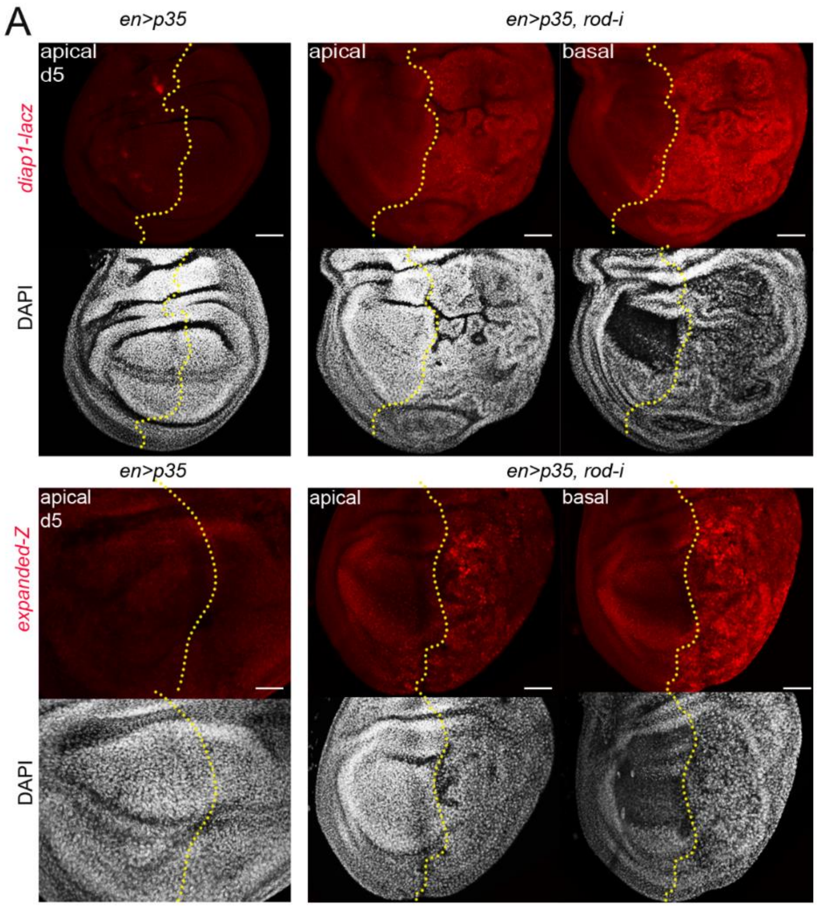


**Figure S3. Yorkie targets are activated in aneuploidy-induced senescent cells** (related to Figure 3)

(**A**) Control wing discs (first column) and wing discs subjected to CIN (remaining columns), stained to visualize *DIAP1-lacZ* (red, **A**), *expanded-lacZ* (red, **A**), and DAPI (white in **A**), and expressing the indicated transgenes in the P compartment under the control of the *en-gal4* driver. Scale bars, 50 µm**.** Discs were dissected 5 days AEL.

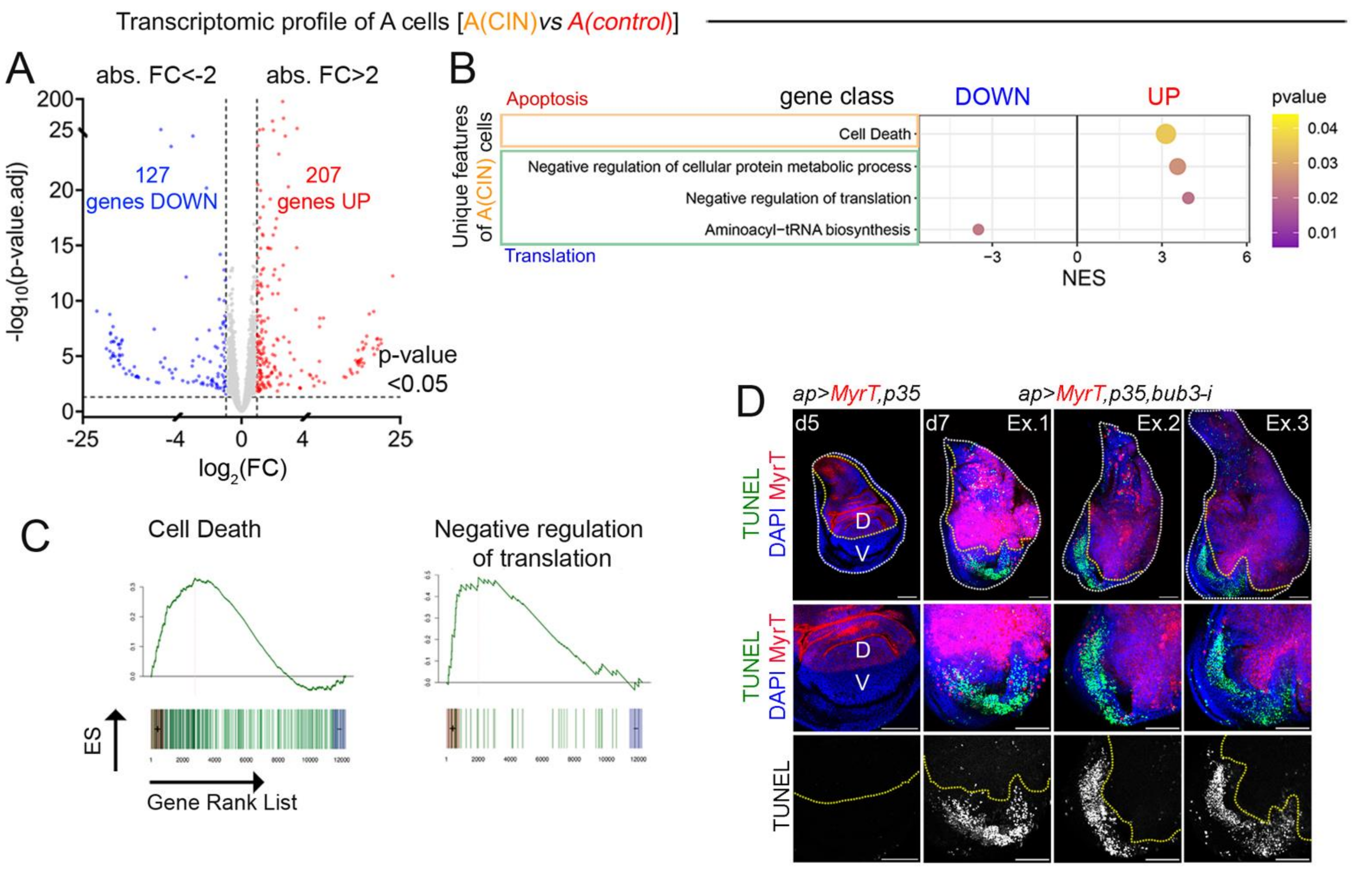


**Figure S4. A non-autonomous effect of CIN tumors on the transcriptomic landscape of the neighboring tissue (**related to Figure 3**)**

**(A)** Volcano plot of all genes that are non-autonomously up- (red dots, FC>2 and p-value<0.05) or down-regulated (blue dots, FC<-2 and p-value<0.05), or not differentially expressed (grey dots, p-value>0.05 or 2>FC>-2) in the A compartment of wing discs bearing CIN tumors in the P compartment when compared to A cells of control discs. **(B)** Bubble plot representing Gene Ontology (GO) enrichment analyses of the A compartment of wing discs bearing CIN tumors in the P compartment when compared to P cells of control discs. Size of the bubble represents the number of measured genes and color-scale represents the associated p-value. **(C)** Mountain plots representing Enrichment Score (ES) upon Gene Set Enrichment Analysis (GSEA) of the A compartment of wing discs bearing CIN tumors in the P compartment when compared to A cells of control discs. GO classes: Cell death - GO0008219, Negative regulation of translation - GO0017148. (**D**) Control wing disc (first column) and three examples of wing discs subjected to CIN (remaining columns), and expressing the indicated transgenes in the dorsal (D) compartment under the control of the *ap-gal4* driver, stained for MyrT (red), TUNEL (green or white) and DAPI (blue). Scale bars, 50 µm**.** Days of dissection after egg laying are as indicated.

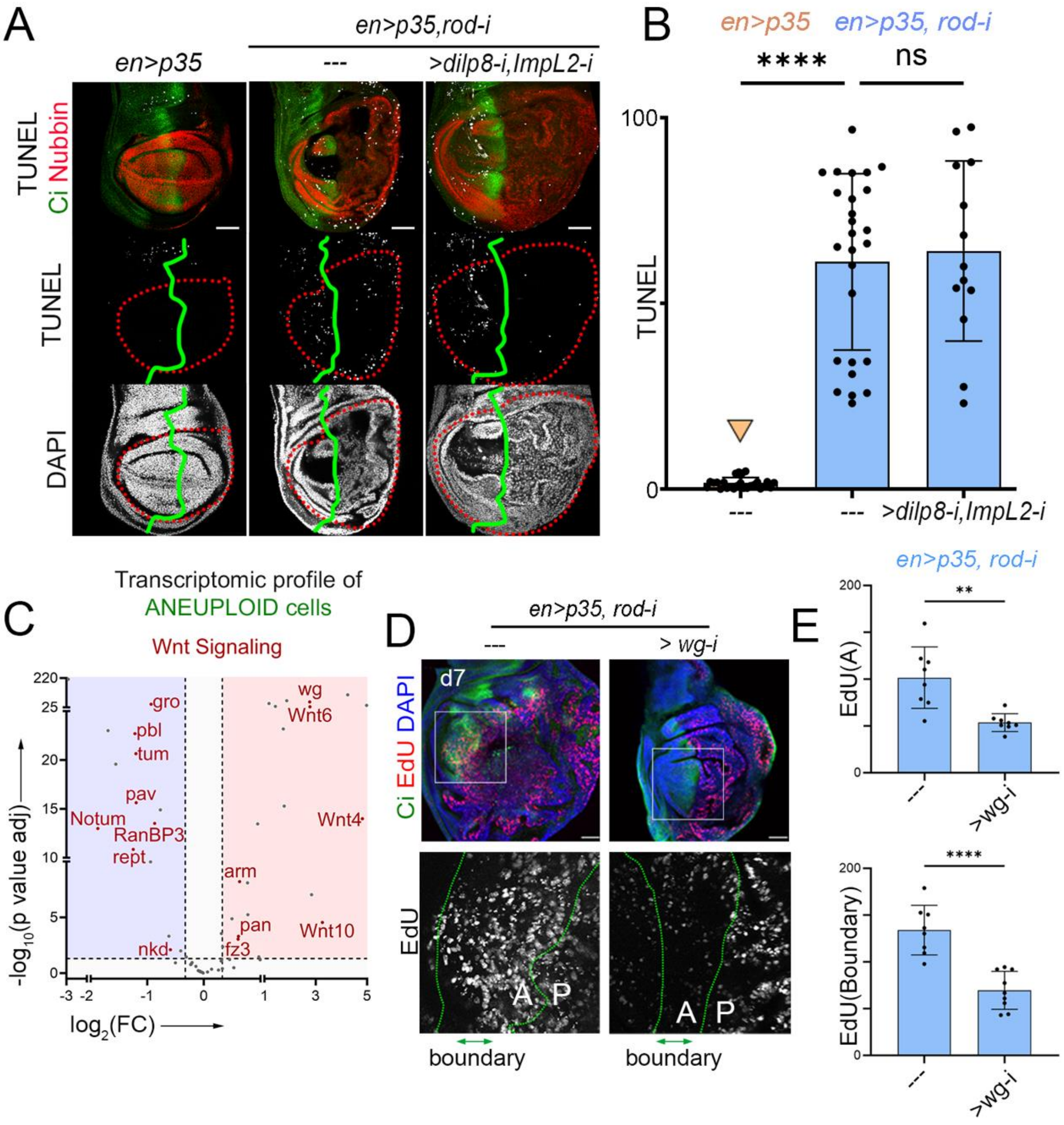


**Figure S5. Contribution of Dilp8/Relaxin, ImpL2/IGFBP7 and Wingless to the non-autonomous effects of CIN tumors on cell death and proliferation** (related to Figure 6)

(**A, D**) Wing discs subjected to CIN, stained to visualize TUNEL (white, **A**), EdU (white, **D**), DAPI (white, white, **A**, or blue, **D**), Nubbin (red, to label the wing pouch cells, **A**), Ci (green, to label the A compartment), and expressing the indicated transgenes in the P compartment under the control of the *en-gal4* driver. Scale bars, 50 µm**.** Discs were dissected 5 days AEL. The boundary between A and P cells is labeled by a green line in **A**. High magnifications of the squared regions are shown in the lower panels in **D**. Note in **D** that the EdU incorporation of the anterior stripe abutting the P compartment in CIN tissues, demarcated by two green lines and double-headed arrow, is not observed upon targeted depletion of *wingless* in the P compartment. (**B**, **E**) Histograms plotting the amount of TUNEL (**B**) or EdU signal (**E**) in the A wing pouch compartments or wing pouch AP boundary (normalized to the size of the corresponding territories) of wing discs expressing the indicated transgenes in the P compartment under the control of the *en-gal4* driver. Ordinary one-way ANOVA was performed for analyzing the data plotted in **B,** Unpaired t test was performed for analyzing the data plotted in **E**; * p<0.05, **p<0.01, ***p<0.001, **** p<0.0001, ns, not significant. **(C)** Volcano plot representing selected genes in 'Wnt Signaling - GO0060070 and GO0090090' classes that are, differentially upregulated (red area, FC>1.3 and p-value<0.05), downregulated (blue area, FC<-1.3 and p-value<0.05), or not differentially expressed (grey area, p-value>0.05 or 1.3>FC>-1.3) in aneuploid cells when compared to P cells of control discs. See also Table S3.

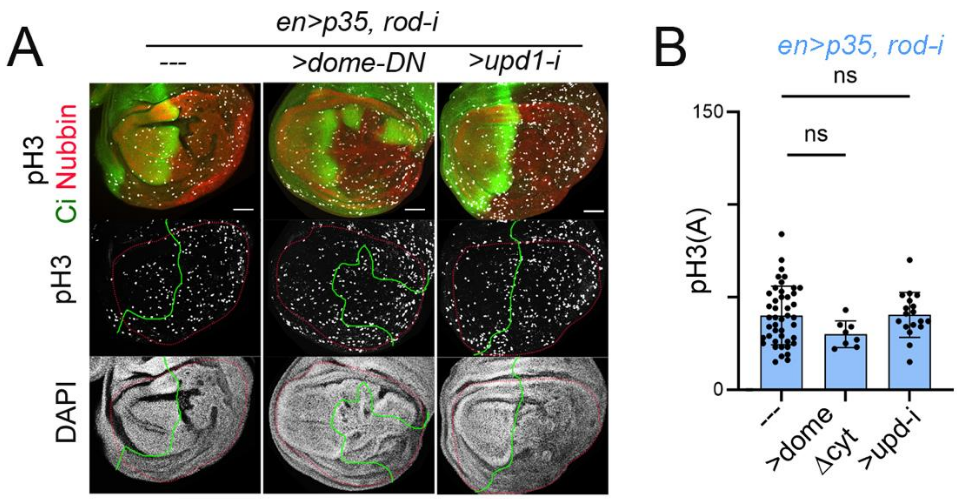


**Figure S6. Contribution of Upd cytiokines to the non-autonomous effects of CIN tumors on cell proliferation** (related to Figure 7)

(**A**) Wing discs subjected to CIN, stained for pH3 (white), Nubbin (red, to label the wing pouch cells) and Ci (green, to label the A compartment), and expressing the indicated transgenes in the P compartment under the control of the *en-gal4* driver. Scale bars, 50 µm**.** Discs were dissected 5cdays AEL. The boundary between A and P cells is labeled by a green line. (**B**) Histogram plotting the amount of pH3 signal in the A wing pouch compartments (normalized to the size of the corresponding territory) of wing discs expressing the indicated transgenes in the P compartment under the control of the *en-gal4* driver. Ordinary one-way ANOVA was performed for analyzing the data plotted. ns, not significant. See also Table S3.

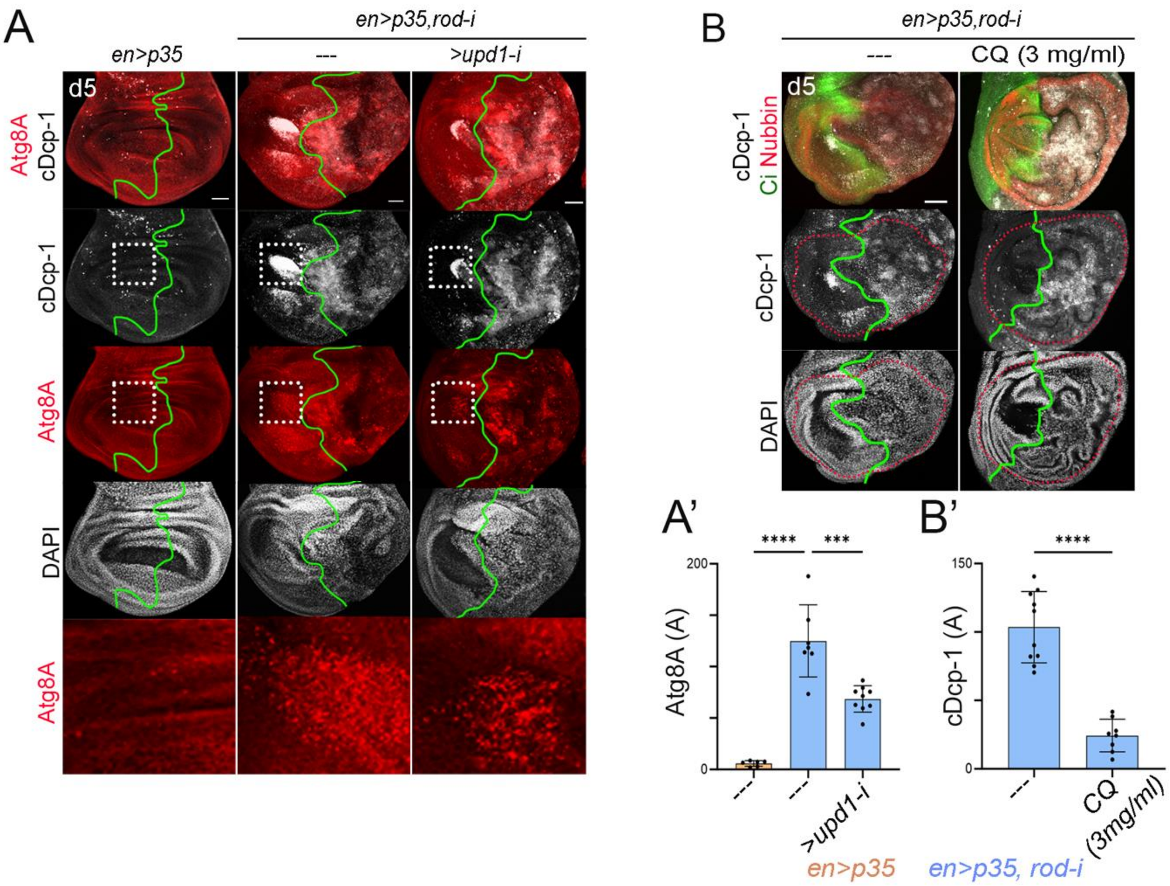


**Figure S7. Cytokine Upd1 contributes induces autophagy cell death** (related to Figure 7)

(**A, B**) Control wing discs (first column in **A**) and wing discs subjected to CIN (remaining columns) stained to visualize cDcp-1 (red and white, **A**; white, **B**), mCherry-Atg8 (red, **A**), Nubbin (red, to label the wing pouch cells, **B**), and Ci (green, to label the A compartment, **B**), and expressing the indicated transgenes in the P compartment under the control of the *en-gal4* driver. Larvae in **B** were subjected to chloroquine (CQ) feeding for 72 h. Scale bars, 50 µm**.** Discs were dissected 5 days AEL. The boundary between A and P cells is labelled by a green or a yellow dashed line. High magnification of the squared regions are shown in the lower panels in **A**. (**A', B'**) Histograms plotting the amount of mCherry-Atg8 signal (**A'**) or cDcp-1 (**B'**) in the A wing pouch compartment (normalized to the size of the corresponding compartment) of wing discs expressing the indicated transgenes in the P compartment under the control of the *en-gal4* driver. Ordinary one-way ANOVA was performed. *p<0.05, **p<0.01, ***p<0.001, **** p<0.0001, ns, not significant. See also Table S3.

**Table S1. Transcriptional profile of CIN, aneuploid cells, and neighboring tissues** (related to Figures 1, 2, 3, 7, S2, S4 and S5)

Excel file containing lists of genes and GO classes of each transcriptional profile with the corresponding fold changes and statistical parameters.

**Table S2. Transcriptional profile of the secretome** (related to Figures 2, and Table 1)

Excel file containing a manually curated list of genes encoding for secreted proteins in the *Drosophila* genome, and its transcriptional profile in aneuploid cells. Fold changes and statistical parameters are included. Table 1 contains the list of top upregulated secreted proteins while comparing the aneuploid cells with control cells, along with the rank of these genes among the rest of the genes.

**Table S3 Summary of n and p-values** (related to Figures 3-7 and S3-S7)

Excel file containing the parameters that have been quantified, the figures where these quantifications are shown, and the statistical details.

**Table S4 Key resources table** (related to Figures 1-7 and S1-S7)

Table containing the list of reagents, chemicals, fly strains and software programs used in this work.

| REAGENT or RESOURCE | SOURCE | IDENTIFIER |
|---|---|---|
| **Antibodies** | | |
| mouse anti-MMP1 (14A3D2)) | Developmental Studies Hybridoma bank | RRID: AB_579782 |
| mouse anti-β-Gal (40.1a) | Developmental Studies Hybridoma bank | RRID: AB_528100 |
| rabbit anti-c-Dcp-1 (9578S) | Cell Signaling Technology | RRID: AB_2721060 |
| rat anti-Ci (2A1) | Developmental Studies Hybridoma bank | RRID: AB_2109711 |
| mouse anti-Nubbin (nub2D4) | Developmental Studies Hybridoma bank | RRID: AB_2722119 |
| rabbit anti-phospho-Histone H3 (pH3) | Cell Signaling | RRID: AB_331535 |
| Alexa fluor 488-conjugated AffiniPure Donkey Anti-Mouse IgG (H+L) | Jackson ImmunoResearch | RRID: AB_2341099 |
| Cy2 AffiniPure Donkey Anti-Rabbit IgG (H+L) | Jackson ImmunoResearch | RRID: AB_2307385 |
| Cy3 AffiniPure Donkey Anti-Mouse IgG (H+L) | Jackson ImmunoResearch | RRID: AB_2315777 |
| Cy3 AffiniPure Donkey Anti-Rabbit IgG (H+L) | Jackson ImmunoResearch | RRID: AB_2307443 |
| Cy5 AffiniPure Donkey Anti-Rat IgG (H+L) | Jackson ImmunoResearch | RRID: AB_2340671 |

| | | |
|---|---|---|
| Donkey anti-Chicken IgY (H+L) Highly Cross Adsorbed Secondary Antibody, Alexa Fluor™ 647 | Invitrogen | RRID: AB_2921074 |
| Chicken Anti-beta Galactosidase Polyclonal Antibody, Unconjugated | Abcam | RRID: AB_307210 |
| **Chemicals, Peptides and Recombinant Proteins** | | |
| Click-iT™ Plus EdU Alexa Fluor™ 647 Imaging Kit | Invitrogen | Code: C10640 |
| DAPI | Sigma Aldrich | Code: 28718-90-3 |
| In Situ Cell Death Detection Kit, Fluorescein (TUNEL) | Roche | Code:11684 795910 |
| Chloroquine | Sigma Aldrich | Code: C6628 |
| **Experimental Models. Organisms/Strains** | | |
| *ap-GAL4* | Bloomington Drosophila Stock Center | RRID: BDSC_3041 |
| *en-GAL4* | Bloomington Drosophila Stock Center | RRID: BDSC_1973 |
| *UAS-bub3RNAi* | Vienna Drosophila Resource Center | RRID: VDRC_21037 |
| *UAS-rodRNAi* | Vienna Drosophila Resource Center | RRID: VDRC_19152 |
| *UAS-p35* | Bloomington Drosophila Stock Center | RRID: BDSC_5072 |
| *UAS-myristoylated-Tomato (myrT)* | Bloomington Drosophila Stock Center | RRID: BDSC_32222 |
| *UAS-Golgi-GFP* | Bloomington Drosophila Stock Center | RRID: BDSC_30902 |
| *UAS-Sec61β-RFP* | Bloomington Drosophila Stock Center | RRID: BDSC_64747 |
| *UAS-CD63-GFP* | Bloomington Drosophila Stock Center | RRID: BDSC_91390 |
| *UAS-bsk-DN* | Bloomington Drosophila Stock Center | RRID: BDSC_6405 |
| *bantam-lacZ* | Bloomington Drosophila Stock Center | RRID: BDSC_10154 |
| *diap1-GFP4.3 (diap1-GFP in the text)* | (1) | - |
| *yorkie-GFP* | (2) | - |
| *Jub-GFP* | Bloomington Drosophila Stock Center | RRID:BDSC_56806 |
| *diap1-lacZ* | (3) | |
| *expanded-lacZ* | (4) | |
| *UAS-YkiRNAi* | Vienna Drosophila Resource Center | RRID: VDRC_40497 |
| *UAS-GC3Ai* | Bloomington Drosophila Stock Center | RRID: BDSC_84343 |

| | | |
|---|---|---|
| *hid-lacZ* | Bloomington Drosophila Stock Center | RRID: BDSC_38010 |
| *MMP1-GFP* | (5) | N/A |
| *Df(H99)* | Bloomington Drosophila Stock Center | RRID:BDSC_1576 |
| *UAS-dilp8RNAi* | Vienna Drosophila Resource Center | RRID: VDRC_102604 |
| *UAS-ImpL2RNAi* | Bloomington Drosophila Stock Center | RRID: BDSC_55855 |
| *UAS-dilp8* | (6) | - |
| *UAS-ImpL2* | (7) | - |
| *dilp8mimicGFP (dilp8-GFP)* | Bloomington Drosophila Stock Center | RRID: BDSC_33079 |
| *impL2mimicGFP (ImpL2-GFP)* | Bloomington Drosophila Stock Center | RRID: BDSC_350068 |
| *UAS-wgRNAi* | Vienna Drosophila Resource Center | RRID: VDRC_104579 |
| *STATGFP10x (II)* | (8) | - |
| *UAS-domeDN (III) (Cyt)* | (9) | - |
| *UAS-domeRNAi* | Vienna Drosophila Resource Center | RRID: VDRC_106071 |
| *hop 27* | Bloomington Drosophila Stock Center | RRID: BDSC_8493 |
| *UAS-upd1RNAi* | Bloomington Drosophila Stock Center | RRID: BDSC_33680 |
| *UAS-upd3RNAi* | Bloomington Drosophila Stock Center | RRID: BDSC_32589 |
| *egr-lacZ* | (10) | - |
| *UAS-grnd-ext (1)* | (11) | - |
| *UAS-grnd-ext (2)* | (11) | - |
| *UAS-grndRNAi* | Vienna Drosophila Resource Center | RRID: VDRC_104538 |
| *UAS-egrRNAi (1)* | Vienna Drosophila Resource Center | RRID: VDRC_108814 |
| *UAS-egrRNAi (2)* | Gifted by Tatsushi Igaki | - |
| *3xmCherry-Atg8a* | (12) | - |
| **Software and Algorithms** | | |
| Fiji | Fiji | https://fiji.sc/ |
| Excel | Microsoft Excel 2021 | N/A |
| GraphPad Prism 10.0.3 | GraphPad | RRID:SCR_002798 |
| Biorender | Biorender | https://www.biorender.com |

**Resource availability**

Lead Contact

**Materials Availability**

The strains generated in the course of this work are freely available to academic researchers through the Lead Contact.
